# Joint effects of climate change and urbanization accelerate plant biodiversity loss across urban green spaces

**DOI:** 10.64898/2026.11.02.763683

**Authors:** Kilian Perrelet, Andreas Dietzel, Lauren M. Cook, Fengge Liu, Joan Casanelles-Abella, Marco Moretti

## Abstract

Urbanization and climate change are rapidly reshaping biodiversity worldwide, yet their combined impacts on ecosystems remain poorly understood. Urban green spaces (UGS) underpin urban biodiversity, but differ widely in their structure, management, and exposure to global change. Here, we examine how these two drivers jointly influence plant communities across five major UGS types (allotments, private gardens, residential green spaces, parks, and ruderal areas) in three socio-culturally distinct Swiss cities. By combining vegetation surveys from 180 sites (60 sites per city) with species-specific thermal niches, high-resolution projections of historical and future urban microclimates, and land-use change scenarios, we trace how climate-driven species loss interacts with the shifting availability of urban habitats to shape biodiversity outcomes by mid-century. We found that UGS types differed markedly in their contributions to biodiversity, with allotments and ruderal areas supporting disproportionately high compositional uniqueness. However, climate change alone was projected to drive substantial plant diversity declines across all UGS types, with 40–80% of species exceeding their thermal niches by mid-century based on climate change scenario and UGS type. Although allotments exhibited relatively greater tolerance due to broader community-level thermal preferences, this advantage was undermined by their rapid decline under urbanization. When considered jointly, climate change and urbanization generated widespread biodiversity losses through complementary mechanisms: climate change drove species declines, while urbanization amplified these losses by disproportionately reducing high-value habitats. These results reveal a consistent trajectory of biodiversity decline across urban landscapes and highlight the need to conserve high-diversity yet shrinking UGS, particularly allotments, while enhancing the ecological value and climate resilience of expanding UGS, such as parks and residential areas, to safeguard future urban biodiversity under global change.

## Introduction

Urbanization and climate change are among the most pervasive drivers of biodiversity loss in the 21st century. By 2050, cities are expected to host around 70% of the global human population, while global temperatures are projected to rise between 1.5–2.5 °C compared to preindustrial era (IPCC 2023). These rapid transformations are already reshaping ecosystems worldwide (Thomas et al. 2004, Grimm et al. 2008, Simkin et al. 2022), and emerging evidence suggests that their impacts are not simply additive: the interaction between urbanization and climate warming can amplify threats to biodiversity beyond what either driver would cause alone (Urban et al. 2024). Climate change contributes to warming at the regional scale, whereas urbanization alters the local microclimate by increasing heat storage in impervious surfaces, such as asphalt, and reducing airflow (Zhao et al. 2021, Urban et al. 2024). As a result, urban microclimates are typically warmer than surrounding areas by an average of 1.6–2.4 °C, depending on the time of day (Varquez and Kanda 2018). As urban areas continue to expand and the climate warms, understanding the combined effects of urban growth and climate change on biodiversity has become a pressing scientific and societal challenge, although these two drivers are rarely considered together in practice (Esperon-Rodriguez et al. 2022, Urban et al. 2024, Dietzel et al. 2025).

Within urban landscapes, species persist primarily in urban green spaces (UGS), which range from intensively managed private gardens and allotments to semi-natural parks and ruderal areas (Swan et al. 2011, Aronson et al. 2017, Fournier et al. 2020). These spaces differ in vegetation structure, composition, and management, generating distinct ecological conditions (Zhao et al. 2010, Gong et al. 2013, Threlfall et al. 2016). Allotments and private gardens often harbor species-rich, cultivated, and non-native (i.e., outside the regional species pool) species, whereas ruderal areas and parks are poorer, often dominated by opportunistic weeds and lawns and trees, respectively (Threlfall et al. 2016; Ruile et al., 2026). These varying plant communities, in turn, shape broader urban biodiversity, influencing the diversity and abundance of bacteria, fungi, invertebrates, and vertebrates (Threlfall et al. 2017; Wagstaffe and Dennis 2025). Plant origin can further modulate these effects, as non-native species may fail to provide key resources due to trait mismatches or the absence of co-evolutionary interactions (Harrison and Winfree 2015, Mata et al. 2021, Tartaglia and Aronson 2024). Collectively, the diversity and composition of plants across UGS types form a mosaic-like foundation of habitats that sustains urban biodiversity (Aronson et al. 2017, Alberti et al. 2020).

Yet this foundation is increasingly threatened. Climate change is projected to push many species beyond their climatic tolerances, with models predicting that around 70% of tree and shrub species in urban forests worldwide risk exceeding their thermal niche by 2050 (Esperon-Rodriguez et al. 2022). This risk has raised the question of whether non-native species, particularly those originating from warmer regions, could thus provide suitable planting options for climate adaptation (Russo et al. 2025). Despite these insights, most of the evidence focuses on a single UGS type—typically forests (Esperon-Rodriguez et al. 2022)—leaving responses of other green space types poorly understood. Furthermore, existing research has largely examined urban growth and climate change separately, overlooking their joint effects (Dietzel et al. 2025). For example, allotments and private gardens, often enriched with warm-adapted non-native species, may act as refugia under future climates (Van der Veken et al. 2008, Haeuser et al. 2019), but they are simultaneously vulnerable to urban densification (Lin et al. 2015, Spilková and Vágner 2016). Lastly, commonly used mesoclimate data (e.g., 30 arc seconds resolution) fail to capture urban microclimates accurately, thereby underestimating climatic risk to species and the loss of suitable habitats within cities due to overwarming (Dietzel et al. 2025). If UGS are to maintain their role as the mosaic underpinning urban biodiversity, efforts must be made to ensure diverse green spaces can withstand the interacting effects of climate change and urban growth.

Here, we assess the unique and combined impacts of urban growth and climate change on urban vegetation across three Swiss cities representing distinct socio-cultural contexts. We integrate vegetation surveys from 180 sites across five UGS types (allotment lots, private gardens, residential areas, parks, and ruderal areas) with high-resolution projections of future urban microclimates and thermal preferences across 970 species. We predict shifts in native and non-native community composition under mid-21st-century climate scenarios and evaluate how changes in UGS availability, driven by urban growth, may interact with climate change to reshape urban plant assemblages. Our approach helps assess how urban biodiversity may persist or decline under interacting global change drivers and offers guidance for designing resilient urban ecosystems.

## Methods

The goal of this study was to assess how UGS support plant biodiversity and how their future contribution may be affected by climate change and urban growth. We first quantified biodiversity across UGS types and characterized species’ thermal niches using global occurrence data (Fig. 1). We then combined these thermal niches with high-resolution microclimate projections for three Swiss cities to assess potential climate-driven species losses. In parallel, we linked historical and current UGS availability to population trends and projected future habitat changes under alternative population growth scenarios (Fig. 1). These projections were then translated into biodiversity changes through simulations of site gains and losses using accumulation curves. Finally, we combined species’ thermal niches and simulations of site gains and losses to assess climate- and land-use-driven combined impacts on future urban plant biodiversity (Fig. 1).

**Figure 1:**
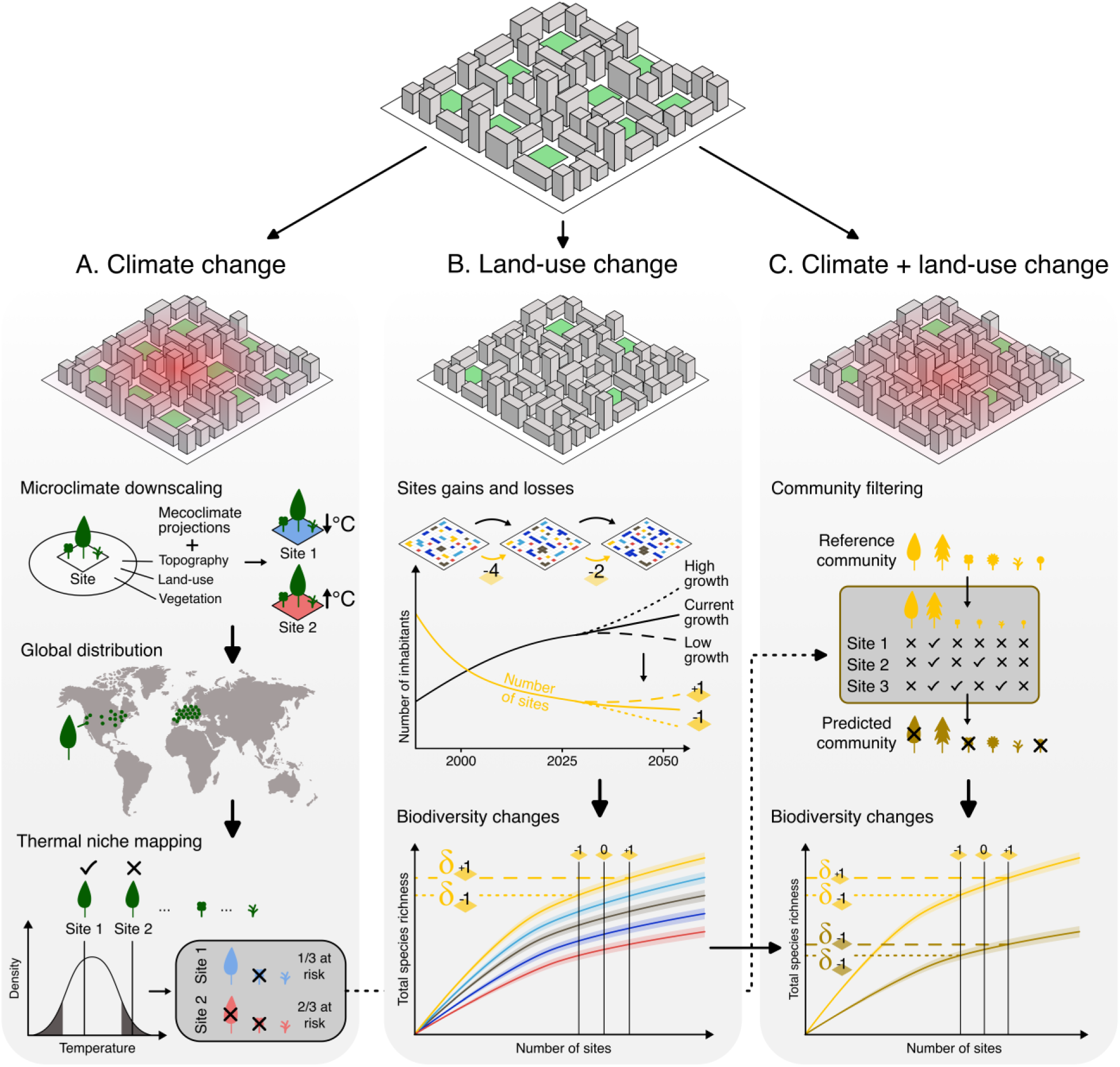
Analytical framework for quantifying the independent (A, B) and combined effects (C) of climate and land-use change on urban plant biodiversity. A: Climate change impacts were estimated by comparing downscaled future microclimates with species’ global thermal niches. B: Land-use change impacts were projected from historical urban green space cover dynamics under alternative population growth scenarios. Deltas indicate examples of biodiversity changes associated with (δ_+1_) or losses (δ_-1_) of sites. C: Combined climate and land-use change impacts were assessed by simulating future communities in which species were first filtered based on expected thermal persistence at each site, followed by land-use projections to estimate biodiversity responses at the landscape-level.

### Species data collection

We used plant community data from Casanelles-Abella et al. (2026) collected across 180 sites in three Swiss cities: Zurich, Geneva, and Lugano (60 sites per city). These cities represent the largest urban centers of the German-, French-, and Italian-speaking regions of Switzerland and were selected to capture both climatic variation—ranging from the cooler, wetter Swiss Plateau (Zurich, Geneva) to the warmer, Mediterranean-influenced Southern Alps (Lugano)—and marked socio-cultural differences (Götz et al. 2018). Within each city, vegetation was surveyed across five UGS types: allotment lots, private gardens, residential green areas (henceforth, “real estate”), parks, and ruderal areas, defined as mostly publicly owned dry grasslands characterized by low overall management intensity but targeted invasive species control (12 sites per type). Sites were selected to represent variability along two orthogonal axes: urban densification and habitat complexity, ensuring wide environmental coverage across the three cities (Fig. 1). Urban densification was measured as the proportion of impervious surface within a 500 m radius, and habitat complexity was quantified based on lawn cover and the number of habitat types present (e.g., meadow, shrubland, trees, flower beds, vegetable beds, wetland). Further details on site selection are provided in Casanelles-Abella et al. (2026).

Vegetation surveys were conducted in May–June 2023 using standardized 100 m^2^ quadrats, with quadrat placement adapted to the specific features of each UGS type. Specifically, quadrats were placed as to capture as much as possible the complexity of each site: for allotments, this represents more or less the size of the lot; for ruderals, the quadrat was placed randomly in the site; for private gardens, the quadrat was subdivided into two 50 m^2^ subquadrats (one in the center of the garden and a second at the edge); for parks and real estates, quadrats were split proportionally to the coverage of the main habitat types of the site (that is, lawn, meadow, ruderal, woodland, shrubland, flowerbed; for details, see Ruile et al. 2026). All plant species were identified to species level by expert botanists with taxonomic treatment following the Swiss Flora checklist (Juillerat et al. 2017) and World Flora Online (WFO 2025). We then classified species as native or non-native following Casanelles-Abella et al. (2021) and as herbs, lianas, shrubs, or trees based on their growth form, following Grothe et al. (2026).

### Biodiversity patterns

To get a first assessment of community richness and uniqueness across UGS types, we quantified species richness and Local Contribution to Beta Diversity (LCBD, Legendre and De Cáceres 2013; 999 permutations) for each site. Differences among UGS types were tested using (generalized) linear mixed-effects models with UGS type as a fixed effect and city as a random intercept (negative binomial distribution for species richness, Gaussian distribution for LCBD). Community composition was assessed using principal coordinate analysis (PCoA) based on Bray–Curtis dissimilarity, with differences among UGS types tested via PERMANOVA (999 permutations).

### Thermal niche characterization

To determine the global thermal niche of each species, we retrieved data on the global distribution of each species from the Global Biodiversity Information Facility (GBIF), restricting our query to observations recorded after 1970, with a maximum coordinate uncertainty of 500 m, and excluding fossil, preserved, and literature-based records, following Esperon-Rodriguez et al. (2022). Species with fewer than 500 global occurrence records were excluded from further analyses. Climatic conditions at occurrence points were extracted from CHELSA V2 (Brun et al. 2022) for Mean Annual Temperature (MAT; bio1) and Mean Temperature of the Warmest Quarter (MTWQ; bio5). For each species, we defined its thermal niche as the temperature values across 95% of its occurrences, assuming that occurrences falling outside the 95% quantile range represent conditions beyond the species’ typical thermal niche, indicating potentially vulnerable populations (Esperon-Rodriguez et al. 2022; Fig. 1). For each species, we then calculated its mean and breadth of MAT and MTWQ, with breadth calculated as the difference between the upper and lower bounds.

To compare central tendencies and variability in thermal preferences between native and non-native species, we compared thermal niche properties (mean and breadth for MAT and MTWQ) between native and non-native species using linear models. To scale these differences to the community level, we calculated community-level thermal niche breadth (for MAT and MTWQ) as the mean MAT and MTWQ breadth, respectively, based on species’ presence-absence in each community. We then tested whether community-level thermal niche breadth varied across UGS types using linear mixed-effects models, including UGS type as a fixed effect and city as a random intercept. We also included species richness as an additional covariate to account for its potential effect on such community-level averages, although the results remained qualitatively unchanged. To assess whether patterns differed by origin, analyses were conducted separately for native and non-native species.

### Microclimate modelling and climate change projections

To assess whether climate change will push species beyond their thermal niches, we accounted for the mismatch between coarse-resolution climate data and fine-scale urban thermal heterogeneity. Although global climate datasets (e.g., CHELSA) are increasingly precise, they still fail to capture urban microclimates due to strong spatial variation in land surface temperature driven by heat-retaining surfaces such as buildings and asphalt (Urban et al. 2024).

To address this, we followed the downscaling approach of Dietzel et al. (2025), integrating mesoclimatic data from CHELSA V2 (Brun et al. 2022) with land-use and remote sensing predictors to generate high-resolution (25 m) microclimate maps for all three cities. Specifically, we implemented a statistical model incorporating topography, impervious surface fraction, NDVI within a 100 m buffer, maximum canopy height, and proximity to water to estimate fine-scale spatial variation in bioclimatic conditions across each city. The model was calibrated in Zurich using air temperature data from a sensor network spread throughout the city and reliably captured observed microclimatic variation (R^2^ = 0.70; see Dietzel et al., 2025). The fitted predictor– response relationships were then transferred to Geneva and Lugano, assuming that the dominant drivers of overwarming are consistent across Swiss cities. Rather than directly predicting absolute temperatures, the model quantified spatial deviations in annual mean temperature (MAT; bio1) and maximum temperature of the warmest quarter (MTWQ; bio5) relative to regional background climate conditions at rural reference weather stations also used by CHELSA. These spatial offsets were subsequently combined with CHELSA projections at the reference stations to generate high-resolution maps of historical (1981–2010) and mid-century (2041–2070) bioclimatic conditions, with the latter selected to align with the temporal framework of land-use projections (see *Land-use change modeling*). Future microclimatic conditions were projected under SSP3-7.0 and SSP5-8.5 scenarios (hereafter “medium” and “high” emissions) and for four climate models (GCMs MPI-ESM1-2-HR, UKESM1-0-LL, and GFDL-ESM4; henceforth “climate model”). Details on microclimate modeling are available in *Supplementary Materials I*.

Species persistence under climate change was then evaluated by comparing, for each climate model, projected site-level temperatures (averaged MAT and MTWQ within each UGS, respectively) with species’ thermal niches. Species were considered persistent only if both projected site-level MAT and MTWQ variables remained within the central 95% of their global climatic distributions. This approach provided site-level estimates of the proportion of potentially vulnerable species, which were compared among UGS types using generalized linear mixed models (beta-binomial distribution). Models included UGS type, emission scenario, and their interaction as fixed effects, and city and climate model as random effects to account for variation among cities and climate projections, and were fitted independently for native and non-native species. Alternative models treating cities and climate projections as fixed effects or including species origin as a fixed effect yielded similar results but higher AIC values and were therefore discarded. We also compared whether species differed in their thermal average or breath based on their origin (native vs. non-native) and growth forms (herbs, lianas, shrubs, and trees) using Wilcoxon and Tukey HSD post-hoc tests.

Although land-use change associated with urbanization is expected to further influence microclimates, spatially explicit future land-use maps are lacking for all three cities, and we thus did not explicitly incorporate the effect of future land-use patterns into microclimate projections. Given ongoing densification and increasing impervious surface cover in Swiss cities (Bosch et al. 2020), which are likely to intensify urban heat island effects (Varquez and Kanda 2018, Cabral et al. 2024), our estimates thus likely represent conservative projections of future temperatures.

### Land-use change modeling

To quantify land-use dynamics, we used Swiss-wide land-use maps providing nationwide land-cover information at 100 m resolution across five time periods (1979–85, 1992–97, 2004–09, 2013–18, and 2020–25; Swiss Federal Statistical Office 2026). For each city and period, we quantified the proportional cover of different UGS types (allotments, private gardens, parks, real estate, and ruderal areas) and related temporal changes in UGS cover to demographic trends using linear mixed-effects models, with population size as a fixed effect and city as a random intercept. This allowed us to account for differences in baseline UGS cover among cities while estimating an overall population–UGS relationship, rather than separate city-specific trends, given the limited number of observations per city. These models were then used to project future UGS availability under three urban growth scenarios (low, current, and high), based on canton-specific population projections (2025–2055; Swiss Federal Statistical Office 2025). Because population projections were only available at the cantonal level and lacked spatial information on future development patterns, spatially explicit predictions of UGS distribution were not possible. Instead, we assessed land-use impacts on biodiversity using a probabilistic framework that accounts for uncertainty in the location of future habitat changes, particularly the creation of new UGS. Although this approach does not capture spatial configuration or local microclimatic variation, it provides a standardized basis for estimating biodiversity responses to changes in the number of suitable sites. As changes were assessed within boundaries defined by population density around cities (see Casanelles-Abella et al. 2026), we define the observed land-use changes as urban growth, encompassing both sprawl and densification.

Projected changes in UGS cover were translated into probabilities of site addition or removal and propagated through 1000 stochastic landscape simulations per climate model and city, generating alternative landscapes with varying numbers of sites for each UGS type, city, and scenario. For each simulated landscape, biodiversity responses were estimated using bootstrapped rarefaction and extrapolation curves, which quantify asymptotic gamma diversity across different numbers of sites (Fig. S1). To account for differences in species pools among UGS types and cities, we generated a separate accumulation curve for each UGS type and city. Changes in site numbers were then applied to the corresponding accumulation curves to estimate changes in city-level species richness under each urban growth scenario.

### Joint effects of changes in climate and land-use

To assess the joint impacts of climate change and urban growth on city-level biodiversity, we combined changes in UGS availability with climate-driven changes in species persistence. For each urban growth scenario, we simulated 1000 future landscapes by randomly removing or adding sites of each UGS type according to projected changes in their availability. We then generated species accumulation curves separately for each city, UGS type, emission scenario, and climate model. For each future climate projection and climate model, we retained only species whose MAT and MTWQ remained within their thermal niches at a given site, thereby generating climate- and model-specific estimates of species accumulation. We evaluated each simulated future landscape by matching its predicted number of sites to the corresponding climate- and model-specific accumulation curve, estimating the expected gamma diversity under that combination of urban growth and climate conditions.

## Results

This study examined the individual and combined effects of climate change and urban growth on urban plant biodiversity across three cities representing diverse socio-cultural contexts. In total, 1,082 taxa were recorded, including 596 in Geneva, 520 in Lugano, and 719 in Zurich. Although patterns of species richness differed among the cities, consistent variation was observed across UGS types (Fig. 2). Allotments, gardens, and real estate supported similar numbers of native species, whereas parks and, particularly, ruderal areas harbored fewer native species (GLMM; *p_park_* = 0.004, *p_ruderal_* < 0.0001; Fig. S2). Contrasts were stronger for non-native species, with allotments being consistently the richest, followed by gardens, real estate, parks, and ruderal areas (GLMM; *p_all_* < 0.0001; Fig. 2, S2). Despite broadly similar proportions of growth forms across UGS types (Fig. S3), community composition differed markedly. Native plant communities in allotments and ruderal areas were more compositionally unique than those in other UGS types (GLMM; *p_ruderal_* = 0.02; *p_all_* < 0.0001; Fig. S4). This pattern was reflected in the PCoA, where allotments and ruderal areas formed distinct clusters, while gardens, parks, and real estate overlapped (PERMANOVA; *p < 0.001*; Fig. S5). In contrast, non-native communities showed similar levels of uniqueness across UGS types, with only ruderal areas contributing slightly less than the others (GLMM; *p_ruderal_* = 0.009; Fig. S4).

**Figure 2:**
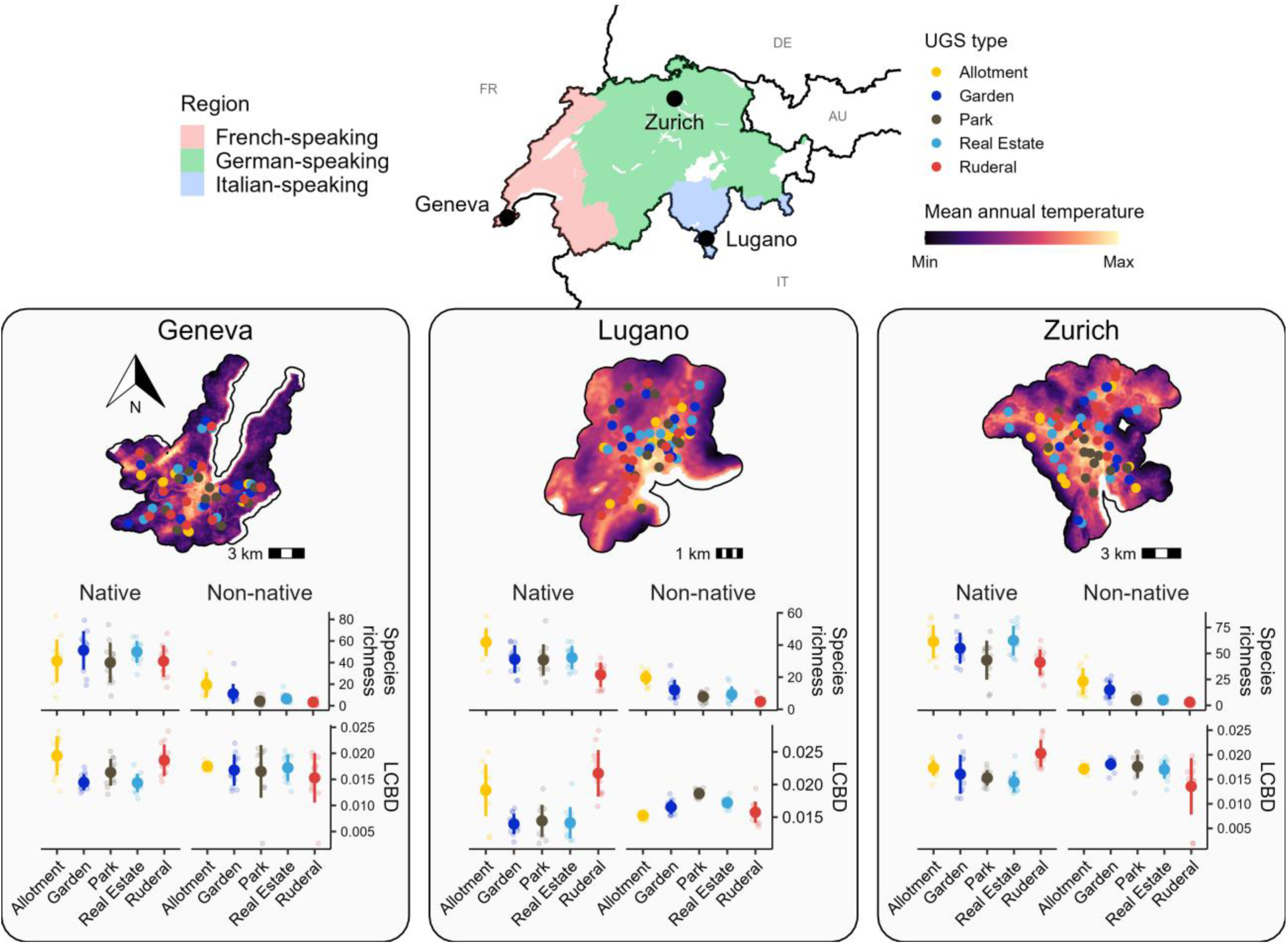
Spatial distribution and biodiversity patterns of urban green spaces (UGS) across Geneva, Lugano, and Zurich. Top panels show city-specific microclimate patterns (min–max scaled within each city; GFDL-ESM4 model), with points indicating sampled UGS locations colored by UGS type. The inset map shows the locations of the three cities within Switzerland. Middle and bottom panels display site-level species richness and local contributions to beta diversity (LCBD) for native and non-native plant assemblages across UGS types. Small points represent individual sites, while larger points and error bars indicate mean ± standard deviation.

UGS types also differed in their vulnerability to climate change (Fig. 3). Already under historical climatic conditions, 241 species (22% of all species, of which 72.2% are native, 27.8% non-native), especially in ruderal areas (80 species, 90% native, 10% non-native), occurred beyond their thermal limits. However, under medium- and high-emission scenarios, 58–63% of native species on average were projected to be pushed beyond their thermal limits across most UGS types and climate models, with species in allotments showing comparatively lower losses (43.2%; Fig. 3; see Table S1–2 for pairwise comparisons between UGS types and emission scenarios). Non-native species exhibited similar but slightly weaker declines (48–59%), again with allotments remaining relatively buffered (30%; Fig. 3). These patterns were broadly consistent across cities but were most pronounced in Lugano, where up to 71.2% of species in ruderal areas were expected to exceed their thermal limits (Fig. S6). These projections were mirrored in differences in thermal niche breadth, with plant communities in allotments consistently exhibiting broader MAT and MTWQ niches than other UGS types (Fig. S7). Notably, this pattern persisted regardless of species origin (i.e., native vs. non-native) and growth form (Fig. S8), despite clear differences in thermal preferences between native and non-native species (Fig. S9) and among herbs, lianas, shrubs, and trees (Fig. S10).

**Figure 3:**
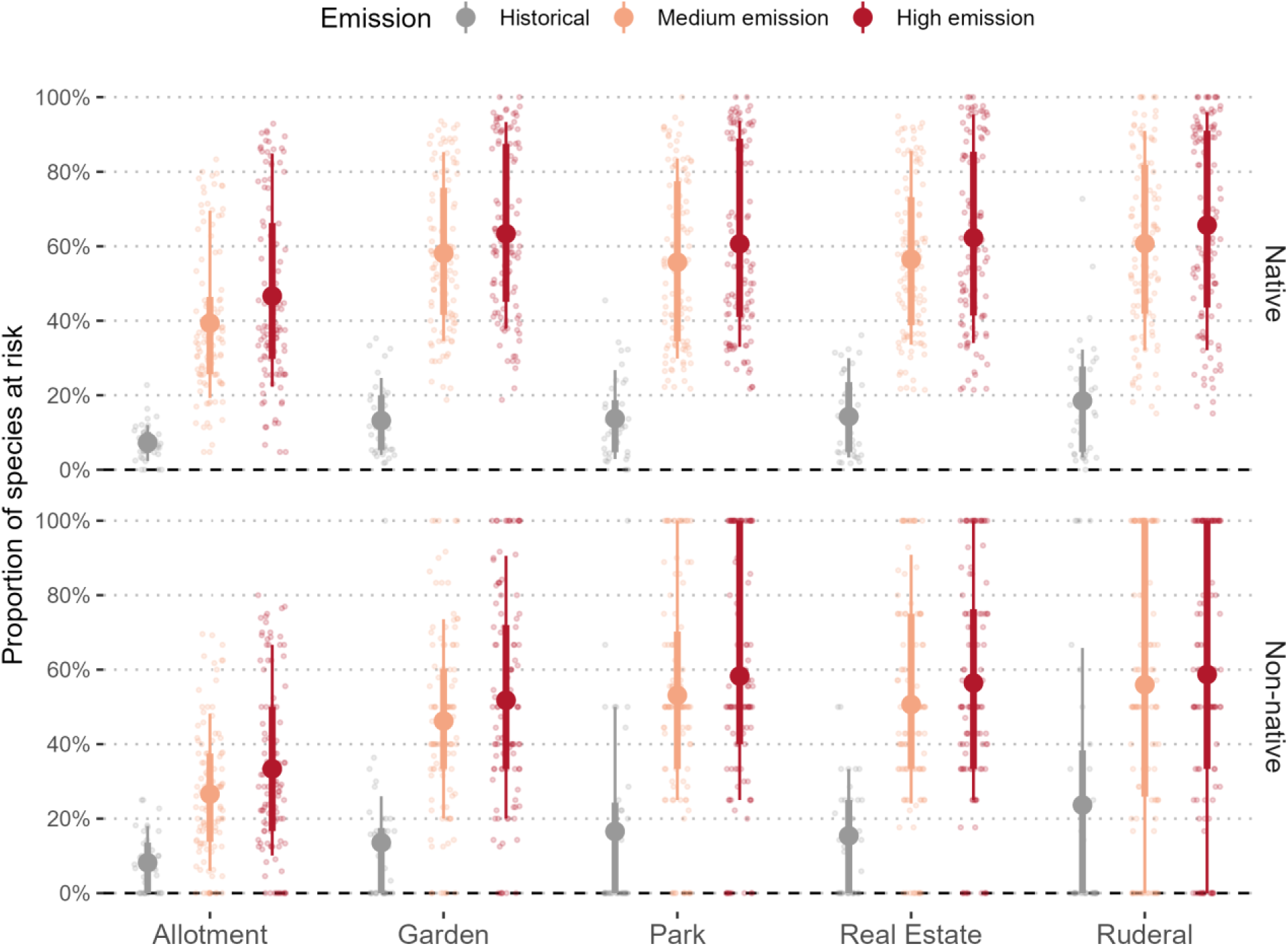
Proportion of native (top) and non-native (bottom) species at risk, defined as occurring beyond their thermal niches, across urban green space (UGS) types under historical and projected future climates. Future projections represent mid-century (2041–2070) conditions under medium-emission (SSP3–7.0) and high-emission (SSP5–8.5) scenarios. Each point represents a site-level estimate for a specific global climate model (MPI-ESM1-2-HR, UKESM1-0-LL, GFDL-ESM4). Large points indicate means, with thick and thin intervals representing 50% and 75% quantile ranges, respectively. Colors denote climate scenarios: historical (grey), medium-emission (light orange), and high-emission (dark red). The dashed horizontal line indicates complete persistence (0% of species at risk). Pairwise comparison coefficients and significance are reported in Tables S1– 2.

Despite the relative climatic tolerance of the communities they harbor, allotment sites were highly vulnerable to urban growth (Fig. 4). Although they currently cover just 0.5–2.9% of urban area across cities (1.4% on average), they experienced the steepest relative decline over the past four decades, shrinking from 1.8% to 1.4% of land cover (21.6% decline; Table S1). Private gardens cover declined from 20.9% to 19.9% (5% decline), whereas parks and real estate areas expanded from 3.3% to 3.8% and 15.7% to 18.8%, respectively (14.4% and 20% increases; Fig. 4, Table S3). These shifts were significantly associated with population growth, particularly for allotments and real estate development, and were most pronounced in Zurich and Geneva (Fig. S11; Table S4). If current growth continues, our projections to 2055 suggest that allotments and gardens will shrink to 1.0% and 19.7% of urban area (further 18% and 5% decline, respectively; Fig. 4). Parks and real estate green spaces, in contrast, are expected to continue expanding (+4.6% and +5.8%, respectively; Table S3), based on linear regressions (Fig. 4). Ruderal areas showed no consistent trend and were not significantly associated with population growth (*p* > 0.05).

**Figure 4:**
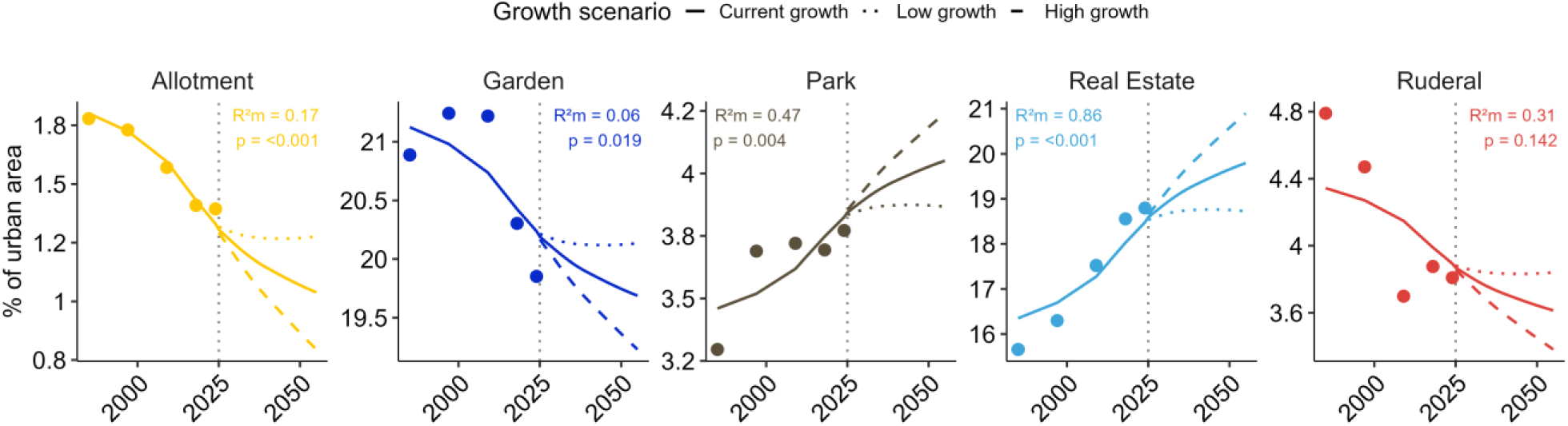
Historical and projected trends in the proportion of UGS types (color-coded) as a function of human population growth. Points show observed historical values averaged across cities, while solid lines represent fitted model relationships. Projections beyond 2025 are shown under three population growth scenarios: current growth (solid), high growth (dashed), and low growth (dotted). Annotations report marginal (m) and conditional (c) R^2^ values, as well as the significance of the human population covariate. Urban areas were delineated based on population density surrounding cities (Casanelles-Abella et al. 2026).

These changes in land cover translated into corresponding gains and losses in biodiversity (Fig. 5, Fig. S12). Under current and high-growth scenarios, gradually disappearing gardens and, especially, allotments were projected to lose 2.3–7.5% and 8.7–25.3% of their current species richness on average across cities, respectively, whereas expanding parks and real estate green spaces showed gains of 2–7.2% and 2.3–6.8%, respectively (Fig. 5, Fig. S12). While the effects of urban growth alone were limited and varied among UGS types, the combination of climate and urban pressures resulted in substantial biodiversity declines across all UGS types (Fig. 5). In gardens, projected climate change alone was projected to push 59.7% of species beyond their thermal limits on average across emission scenarios and climate models (Fig. 3), whereas incorporating urban growth effects increased total projected losses to 67.2% under high-growth scenarios (Fig. 5). In allotments, species communities’ greater resistance to climate change (Fig. 3) was offset by higher sensitivities to urban growth (Fig. 4), resulting in losses of up to 65.3% under combined climate and urban pressures, comparable to those projected across other UGS types (64–70.5%; Fig. 5). Similarly, while the expansion of parks and real estate green spaces locally increased species richness (Fig. S12), these gains were consistently outweighed by climate-driven losses, leading to average net declines of 52.2–66.2% depending on urban growth scenario (Fig. 5). Declines in non-native species broadly followed those of native taxa but were slightly less pronounced. Nevertheless, even small differences in persistence between native and non-native species were projected to drive substantial compositional shifts, particularly in ruderal areas, where non-native species increased by 49.6–78.4% relative to current levels, compared with 10.3–27.7% in other UGS types (Fig. S13).

**Figure 5:**
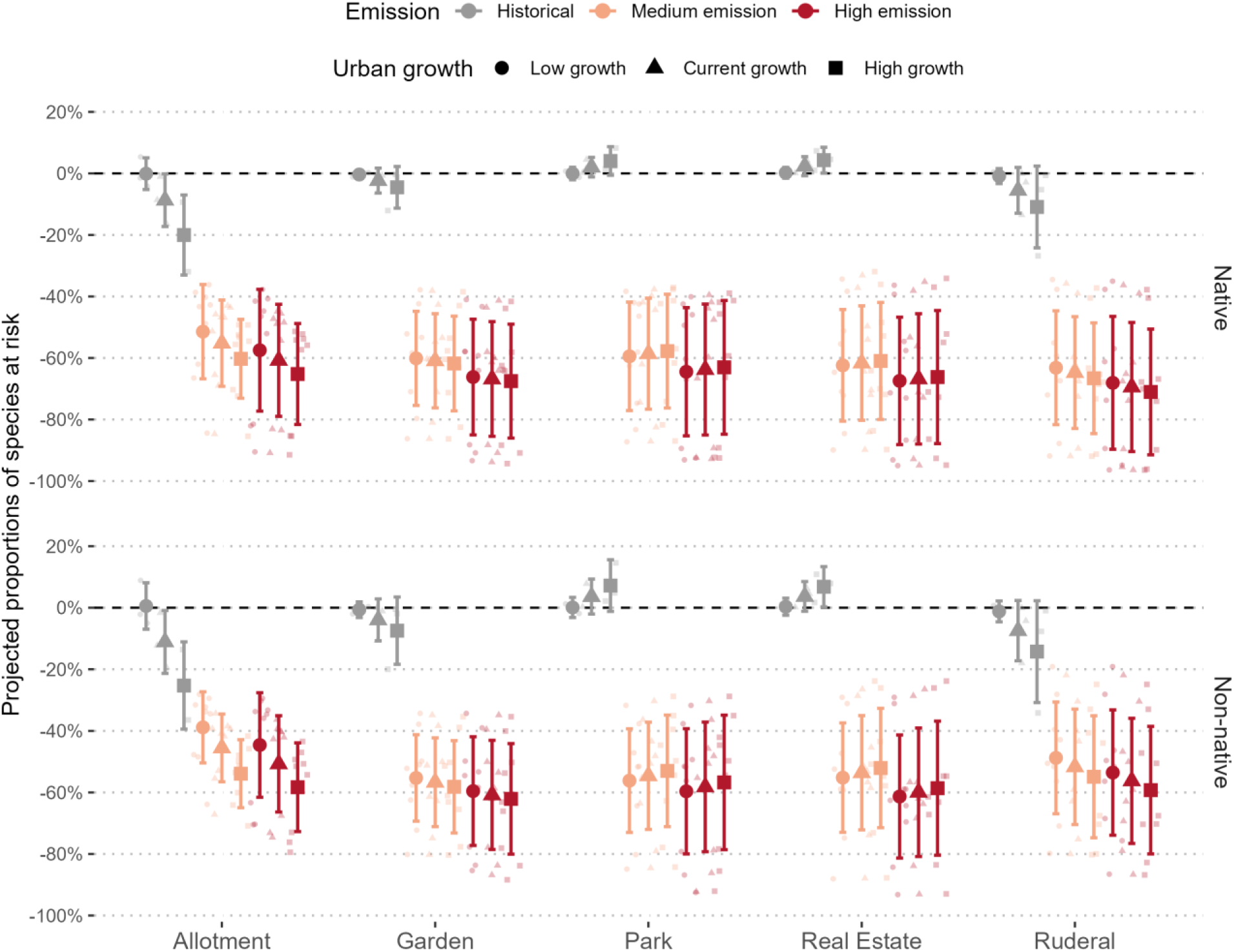
Simulated proportional changes in city-level proportions of species at risk of native (left) and non-native (right) assemblages across urban green space (UGS) types, relative to current baselines. Each small point represents a city (Zurich, Geneva, Lugano) under a specific climate model (MPI-ESM1-2-HR, UKESM1-0-LL, GFDL-ESM4), with values indicating mean proportional changes in species richness from 1000 simulations of site loss and gain under projected land-use change. Urban growth scenarios (low, current, high) are indicated by point shape and line type, and climate scenarios (historical, medium emission, high emission) by color. Larger points and error bars show means ± standard deviation. The dashed black line represents the current baseline; values under the line indicate losses, while values above the line indicate gains.

## Discussion

Urban plant biodiversity is projected to decline markedly under the combined pressures of climate change and urban growth across cities, UGS types, and both native and non-native assemblages. Climate change emerged as the dominant driver, exposing 54–59% (37.9% in allotments) of native and 43–52% of non-native species to substantial climate risks by mid-century, with up to 81% of species in ruderal areas projected to exceed their thermal limits. Urban growth compounded these losses by reshaping habitat availability, contributing up to 21.7% additional losses in gardens and up to 51.6% in allotments.

UGS types differed markedly in species richness, community composition, and compositional uniqueness (LCBD), underscoring the importance of management and anthropogenic purpose (e.g., food versus shade provision) in shaping urban plant communities (Zhao et al. 2010, Threlfall et al. 2016, Aronson et al. 2017). Allotments, and to a lesser extent ruderal areas, were the main contributors to urban biodiversity, combining high species richness with high distinctiveness despite contrasting ecological dynamics (intensive cultivation versus low intervention; Threlfall et al. 2016; Ruile et al., 2026). Human influence was particularly evident in allotments, which supported many non-native ornamental and food species (Speak et al. 2015), whereas ruderal areas disproportionately contributed to compositional distinctiveness, likely acting as reservoirs of stress-tolerant and often rare species (Bonthoux et al. 2014, Sikorski et al. 2021). These patterns were consistent across cities despite climatic and socio-cultural differences (Casanelles-Abella et al. 2026), suggesting that the ecological roles of UGS types may generalize across urban systems.

Climate change, however, threatens this biodiversity by pushing species beyond their realized thermal niches (Esperon-Rodriguez et al. 2022), with risks further exacerbated by urban microclimatic conditions (Dietzel et al. 2025). We projected an average risk of loss of up to 65% by mid-century across cities and climate models, with similar outcomes under both emission scenarios and consistent with projections for urban animals (Dietzel et al. 2025). Losses were lower in allotments, likely reflecting broader thermal niches shaped by gardening practices, including cultivation by gardeners from warmer regions (Young and Bauer 2022), and the frequent use of widely distributed ornamental and food plants, even among native species (Altman et al. 2022). Nevertheless, many non-native species are projected to approach or exceed their thermal limits, whereas many native species are expected to persist even under high-emission scenarios. Native species should therefore not be assumed to be inherently less suitable under future climates, as thermal niches across species’ global distributions may provide a better guide for climate-resilient planting than species origin alone (Hoyle et al. 2017, Alizadeh and Hitchmough 2018, Russo et al. 2025). These projections should, however, be interpreted in light of water availability and management, which strongly influence plant persistence (Meineke and Frank 2018, Tomatis et al. 2023). In intensively managed UGS, particularly allotments and private gardens, irrigation and reseeding by gardeners and managers can buffer climate impacts and help maintain these habitats as refugia for biodiversity and human well-being (Ayling et al. 2021, Tomatis et al. 2023), especially when groundwater is unavailable (Yin et al. 2026). Yet, warmer summers across the three studied cities, which tend to coincide with drier conditions (Yin et al. 2026), may reduce this buffering capacity, with impacts further compounded by increasing drought frequency and associated water-use restrictions across much of urban Switzerland (Milano et al. 2015, Brunner et al. 2019). Addressing this challenge will require transdisciplinary approaches with engineering, social sciences, and local communities to develop water conservation strategies, including rainwater harvesting, greywater reuse, alternative soil composition to enhance water retention, and adaptive irrigation practices (Misra et al. 2010, Cavadini et al. 2024, Fletcher et al. 2024).

Urban growth further modulated biodiversity loss by reshaping the distribution of UGS types, particularly through declines in allotments and, to a lesser extent, private gardens. These trends, which we linked to population growth, mirrored those observed in cities worldwide (Lin et al. 2015, Spilková and Vágner 2016). Importantly, across UGS types, native species declined slightly more than non-native species, highlighting that even modest differences in persistence can gradually shift community composition towards greater dominance of non-native species, especially if planting practices remain unchanged (Russo et al. 2025). Among all UGS types, allotments were particularly vulnerable. Despite occupying a small proportion of urban land (here, <2% on average across cities), allotments are major and unique contributors to both urban fauna and flora (Baldock et al. 2015, Threlfall et al. 2016, Ruile et al. 2026). Yet, their cover declined by over 20% across the three focal cities in recent decades, representing a disproportionate loss of high-value habitat to more intensive land uses (Spilková and Vágner 2016). Although harbouring relatively climate-tolerant species, allotments became as vulnerable as other UGS types once urban growth was considered, highlighting the importance of evaluating interacting global change drivers (Urban et al. 2024). Given the disproportionate contribution of allotments to urban biodiversity, preserving allotments as multifunctional systems will therefore be critical for supporting biodiversity conservation, climate adaptation, and social benefits (e.g., food production, mental restoration; Cabral et al. 2017, Young et al. 2020, Ayling et al. 2021). Private gardens were also projected to decline, particularly in rapidly densifying cities worldwide (Lin et al. 2015, Herdt and Jonkman 2023), although their large spatial extent meant that biodiversity losses remained relatively modest. However, climate change is expected to exacerbate these losses, highlighting the importance of individual planting and management choices. Such choices, which already contribute to substantial variation in plant community composition and associated biodiversity among gardens (Home et al. 2019), offer opportunities to enhance climate resilience through the selection of drought-tolerant or post-drought resprouting species (Chu et al. 2025).

In contrast, urban growth, particularly through urban sprawl, replaced peri-urban, often agricultural land and increased the area covered by parks and residential green spaces, leading to biodiversity gains with increasing levels of growth. However, their comparatively low species richness and compositional distinctiveness mean that these gains cannot compensate for losses of higher-value habitats, such as allotments, and their often unique fauna and flora. Moreover, climate-driven declines consistently outweighed these potential gains. Given their projected increase, parks and residential green spaces nevertheless represent important opportunities for biodiversity promotion (Aronson et al. 2017, Lerman et al. 2023). Increasing their ecological value and climate resilience will require changes in design and management, including lower maintenance intensity, targeted planting of drought-tolerant or resprouting species, and measures that improve water availability, such as stormwater harvesting (He et al. 2021, Marchin et al. 2022, Chu et al. 2025). Because these spaces are widely used, involving local communities in their codesign (e.g., by integrating food production into ornamental and recreational areas) could also improve public acceptance, strengthen environmental stewardship, and generate benefits for both biodiversity and people (Egerer et al. 2024, Kowarik et al. 2025).

By integrating microclimate and land-use projections, our study highlighted substantial plant biodiversity losses under concurrent global change drivers. However, several limitations should be acknowledged. First, we considered only air temperature, represented at 2 m above ground in CHELSA, which may not fully reflect the microclimatic conditions experienced by plants (Klinges et al. 2026), while overlooking other important constraints, such as water availability and soil conditions (Ibáñez et al. 2014, Meineke and Frank 2018). Even within this simplified framework, projections varied considerably among climate models, highlighting the uncertainty associated with future climate forecasts and their implications for biodiversity (IPCC 2023, Urban et al. 2024). In addition, some species currently occur outside their estimated thermal niches, potentially reflecting extinction debt but also uncertainties in defining urban thermal niches from broad-scale data. Urban populations may already be adapted to warmer conditions (Carfora et al. 2025), while microrefugia can decouple climatic exposure from realized occupancy (Kibria et al. 2026), allowing persistence despite unsuitable conditions during some life stages (e.g., seedling versus adult survival; Ibáñez et al. 2014, Canham and Murphy 2016). Our approach also ignored intraspecific variation, further obscuring the thermal niches of urban populations and potentially explaining why some individuals persisted beyond estimated niche limits. Given the importance of plant nurseries as a source of urban vegetation (Grothe et al. 2026), future studies should consider how the widespread use of clonally propagated plants, with reduced genetic variability, may influence climate risks for urban plants (Vanden Broeck et al. 2018). Second, the absence of spatially explicit population and land-use projections limited our ability to capture future changes in habitat configuration (e.g., connectivity) that influence species persistence and colonization (Vega and Küffer 2021). It also prevented us from accounting for feedbacks between urban growth and microclimate, likely resulting in conservative estimates of future urban warming (Varquez and Kanda 2018, Cabral et al. 2024). Under these constraints, we used species accumulation curves, which provided standardized gamma-diversity estimates despite lacking detailed or spatially explicit site-level information (e.g., patch size or distribution). Future studies should therefore integrate spatially explicit frameworks to better capture habitat configuration effects. Finally, we assumed static species pools (i.e., no invasion) and did not account for climate-driven range shifts that may alter urban assemblages (Lososová et al. 2018, Filazzola et al. 2024), or for compositional changes resulting from deliberate and accidental introductions linked to trade, shifting planting preferences, and policy decisions (Mack and Lonsdale 2001, Bradley et al. 2012). Future studies should thus aim at integrating ecological modelling with spatial planning and socio-ecological processes to better anticipate urban biodiversity futures. Despite these uncertainties, our findings emphasize that safeguarding urban biodiversity will require simultaneously addressing climate change and urban growth and that preserving allotments and improving the biodiversity value and climate tolerance of parks and residential green spaces will be central to designing climate-adaptive cities.

## Conclusion

Our findings underscore the critical importance of considering multiple global change drivers when assessing the future of urban biodiversity. In particular, they highlight the disproportionate role of certain UGS types, especially allotments, in sustaining diverse and unique plant communities, as well as their vulnerability to ongoing urban growth. Conservation strategies should therefore prioritize the protection and integration of such high-value habitats within urban planning, while also enhancing the ecological value and climate tolerance of otherwise expanding urban green spaces (e.g., residential areas) through diversified yet targeted planting of drought-tolerant or resprouting species, reduced maintenance intensity, and sustainable water management. Failure to address both climate change and urban growth in an integrated manner risks substantial and potentially irreversible losses and homogenization of urban biodiversity in the coming decades.

## Data availability

Data and code is available on GitHub (https://github.com/KPerrelet/urban_climate_joint_impact) and Zenodo (https://doi.org/10.5281/zenodo.23889283).

## Author contributions

**KP**: Conceptualization, Data Curation, Formal Analysis, Investigation, Methodology, Visualization, Writing – Original Draft Preparation; **AD**: Conceptualization, Methodology, Writing – Review & Editing; **LMC**: Conceptualization, Funding Acquisition, Writing – Review & Editing; **FL**: Formal Analysis, Methodology, Writing – Review & Editing; **JAC**: Investigation, Writing – Review & Editing; **MM**: Conceptualization, Funding Acquisition, Project Administration, Supervision, Writing – Review & Editing.

## Acknowledgments

We thank all the participating stakeholders for their invaluable contribution to the PAPPUS Project.

## References

Alberti, M., Palkovacs, E. P., Roches, S. D., Meester, L. D., Brans, K. I., Govaert, L., Grimm, N. B., Harris, N. C., Hendry, A. P., Schell, C. J., Szulkin, M., Munshi-South, J., Urban, M. C. and Verrelli, B. C. 2020. The Complexity of Urban Eco-evolutionary Dynamics. - BioScience 70: 772–793.

Alizadeh, B. and Hitchmough, J. 2018. A review of urban landscape adaptation to the challenge of climate change. - International Journal of Climate Change Strategies and Management 11: 178–194.

Altman, A., Shennan, S. and Odling-Smee, J. 2022. Ornamental plant domestication by aesthetics-driven human cultural niche construction. - Trends in Plant Science 27: 124–138.

Aronson, M. F., Lepczyk, C. A., Evans, K. L., Goddard, M. A., Lerman, S. B., MacIvor, J. S., Nilon, C. H. and Vargo, T. 2017. Biodiversity in the city: key challenges for urban green space management. - Frontiers in Ecology and the Environment 15: 189–196.

Ayling, S. M., Phillips, N. and Bunney, S. 2021. Allotments in the Future: Building Resilience to Climate Change through Improved Site Design and Efficient Water Practices. - Water in press.

Baldock, K. C. R., Goddard, M. A., Hicks, D. M., Kunin, W. E., Mitschunas, N., Osgathorpe, L. M., Potts, S. G., Robertson, K. M., Scott, A. V., Stone, G. N., Vaughan, I. P. and Memmott, J. 2015. Where is the UK’s pollinator biodiversity? The importance of urban areas for flower-visiting insects. - Proceedings of the Royal Society B: Biological Sciences 282: 20142849.

Bonthoux, S., Brun, M., Di Pietro, F., Greulich, S. and Bouché-Pillon, S. 2014. How can wastelands promote biodiversity in cities? A review. - Landscape and Urban Planning 132: 79–88.

Bosch, M., Jaligot, R. and Chenal, J. 2020. Spatiotemporal patterns of urbanization in three Swiss urban agglomerations: insights from landscape metrics, growth modes and fractal analysis. - Landscape Ecol 35: 879–891.

Bradley, B. A., Blumenthal, D. M., Early, R., Grosholz, E. D., Lawler, J. J., Miller, L. P., Sorte, C. J., D’Antonio, C. M., Diez, J. M., Dukes, J. S., Ibanez, I. and Olden, J. D. 2012. Global change, global trade, and the next wave of plant invasions. - Frontiers in Ecology and the Environment 10: 20–28.

Brun, P., Zimmermann, N. E., Hari, C., Pellissier, L. and Karger, D. N. 2022. CHELSA-BIOCLIM+ A novel set of global climate-related predictors at kilometre-resolution.

Brunner, M. I., Björnsen Gurung, A., Zappa, M., Zekollari, H., Farinotti, D. and Stähli, M. 2019. Present and future water scarcity in Switzerland: Potential for alleviation through reservoirs and lakes. - Science of The Total Environment 666: 1033–1047.

Cabral, I., Costa, S., Weiland, U. and Bonn, A. 2017. Urban Gardens as Multifunctional Nature-Based Solutions for Societal Goals in a Changing Climate. - In: Kabisch, N. et al. (eds), Nature-Based Solutions to Climate Change Adaptation in Urban Areas: Linkages between Science, Policy and Practice. Springer International Publishing, pp. 237–253.

Cabral, J. S., Mendoza-Ponce, A., da Silva, A. P., Oberpriller, J., Mimet, A., Kieslinger, J., Berger, T., Blechschmidt, J., Brönner, M., Classen, A., Fallert, S., Hartig, F., Hof, C., Hoffmann, M., Knoke, T., Krause, A., Lewerentz, A., Pohle, P., Raeder, U., Rammig, A., Redlich, S., Rubanschi, S., Stetter, C., Weisser, W., Vedder, D., Verburg, P. H. and Zurell, D. 2024. The road to integrate climate change projections with regional land-use–biodiversity models. - People and Nature 6: 1716–1741.

Canham, C. D. and Murphy, L. 2016. The demography of tree species response to climate: seedling recruitment and survival. - Ecosphere 7: e01424.

Carfora, A., Lucibelli, F., Di Lillo, P., Mazzucchiello, S. M., Saccone, G., Salvemini, M., Varone, M., Volpe, G. and Aceto, S. 2025. Genetic responses of plants to urban environmental challenges. - Planta 261: 102.

Casanelles-Abella, J., Frey, D., Müller, S., Aleixo, C., Alós Ortí, M., Deguines, N., Hallikma, T., Laanisto, L., Niinemets, Ü., Pinho, P., Samson, R., Villarroya-Villalba, L. and Moretti, M. 2021. A dataset of the flowering plants (Angiospermae) in urban green areas in five European cities. - Data in Brief 37: 107243.

Casanelles-Abella, J., Ruile, S. R., Bergamini, A., Dädlow, G. M. L., Grothe, V., Fournier, B., Perrelet, K. and Moretti, M. 2026. Social-ecological signatures in plant species composition across Swiss cities and urban green space types. - Plants, People, Planet in press.

Cavadini, G. B., Rodriguez, M., Nguyen, T. and Cook, L. M. 2024. Can blue-green infrastructure counteract the effects of climate change on combined sewer overflows? Study of a Swiss catchment. - Environmental Research Letters 19: 094025.

Chu, L. H.-H., Szota, C., Arndt, S. K. and Farrell, C. 2025. What doesn’t kill you makes you stronger: Using plant traits to inform species selection for naturalistic plantings in hotter and drier climates. - Landscape and Urban Planning 259: 105364.

Dietzel, A., Moretti, M., Perrelet, K. and Cook, L. M. 2025. Urban heat exacerbates climatic risks to urban biodiversity. - npj Urban Sustain 6: 4.

Egerer, M., Karlebowski, S., Conitz, F., Neumann, A. E., Schmack, J. M. and Sturm, U. 2024. In defence of urban community gardens. - People and Nature 6: 367–376.

Esperon-Rodriguez, M., Tjoelker, M. G., Lenoir, J., Baumgartner, J. B., Beaumont, L. J., Nipperess, D. A., Power, S. A., Richard, B., Rymer, P. D. and Gallagher, R. V. 2022. Climate change increases global risk to urban forests. - Nat. Clim. Chang. 12: 950–955.

Filazzola, A., Johnson, M. T. J., Barrett, K., Hayes, S., Shrestha, N., Timms, L. and MacIvor, J. S. 2024. The great urban shift: Climate change is predicted to drive mass species turnover in cities. - PLOS ONE 19: e0299217.

Fletcher, T. D., Burns, M. J., Russell, K. L., Hamel, P., Duchesne, S., Cherqui, F. and Roy, A. H. 2024. Concepts and evolution of urban hydrology. - Nat Rev Earth Environ 5: 789–801.

Fournier, B., Frey, D. and Moretti, M. 2020. The origin of urban communities: From the regional species pool to community assemblages in city. - Journal of Biogeography 47: 615–629.

Gong, C., Chen, J. and Yu, S. 2013. Biotic homogenization and differentiation of the flora in artificial and near-natural habitats across urban green spaces. - Landscape and Urban Planning 120: 158–169.

Götz, F. M., Ebert, T. and Rentfrow, P. J. 2018. Regional Cultures and the Psychological Geography of Switzerland: Person–Environment–Fit in Personality Predicts Subjective Wellbeing. - Front. Psychol. in press.

Grimm, N. B., Faeth, S. H., Golubiewski, N. E., Redman, C. L., Wu, J., Bai, X. and Briggs, J. M. 2008. Global Change and the Ecology of Cities. - Science 319: 756–760.

Grothe, V., Dädlow, G. M. L., Casanelles, J., Ruile, S., Moretti, M. and Fournier, B. 2026. The Human-Imprinted Regional Species Pool: Functional Trait Patterns of Commercial and Established Plants.- Authorea in press.

Haeuser, E., Dawson, W. and van Kleunen, M. 2019. Introduced garden plants are strong competitors of native and alien residents under simulated climate change. - Journal of Ecology 107: 1328–1342.

Harrison, T. and Winfree, R. 2015. Urban drivers of plant-pollinator interactions. - Functional Ecology 29: 879–888.

He, C., Liu, Z., Wu, J., Pan, X., Fang, Z., Li, J. and Bryan, B. A. 2021. Future global urban water scarcity and potential solutions. - Nat Commun 12: 4667.

Herdt, T. and Jonkman, A. R. 2023. The acceptance of density: Conflicts of public and private interests in public debate on urban densification. - Cities 140: 104451.

Home, R., Lewis, O., Bauer, N., Fliessbach, A., Frey, D., Lichtsteiner, S., Moretti, M., Tresch, S., Young, C., Zanetta, A. and Stolze, M. 2019. Effects of garden management practices, by different types of gardeners, on human wellbeing and ecological and soil sustainability in Swiss cities. - Urban Ecosyst 22: 189–199.

Hoyle, H., Hitchmough, J. and Jorgensen, A. 2017. Attractive, climate-adapted and sustainable? Public perception of non-native planting in the designed urban landscape. - Landscape and Urban Planning 164: 49–63.

Ibáñez, B., Ibáñez, I., Gómez-Aparicio, L., Ruiz-Benito, P., García, L. V. and Marañón, T. 2014. Contrasting effects of climate change along life stages of a dominant tree species: the importance of soil–climate interactions. - Diversity and Distributions 20: 872–883.

IPCC 2023. Climate change 2023: synthesis report. Contribution of working groups I, II and III to the sixth assessment report of the intergovernmental panel on climate change.

Juillerat, P., Bäumler, B., Bornand, C., Gygax, A., Jutzi, M., Möhl, A., Nyffeler, R., Sager, L., Santiago, H. and Eggenberg, S. 2017. Flora Helvetica Checklist 2017: der Gefässpflanzen der Schweiz = de la flore vasculaire de la Suisse = della flora vascolare della Svizzera. in press.

Kibria, M. G., Tjoelker, M. G., Esperón-Rodríguez, M., Marchin, R. M., Arndt, S. K., Livesley, S. J. and Rymer, P. D. 2026. Functional traits are better than climate niche approach at predicting the vital growth and mortality rates of urban tree species. - Urban Forestry & Urban Greening 120: 129450.

Klinges, D. H., Muñoz, M. M., Domínguez-Guerrero, S. F., Maclean, I. M. D., Kearney, M. R. and Skelly, D. K. 2026. Matching climate to biological scales. - Trends in Ecology & Evolution 41: 329–343.

Kowarik, I., Fischer, L. K., Haase, D., Kabisch, N., Kleinschroth, F., Konijnendijk, C., Straka, T. M. and von Haaren, C. 2025. Promoting urban biodiversity for the benefit of people and nature. - Nat. Rev. Biodivers. 1: 214–232.

Legendre, P. and De Cáceres, M. 2013. Beta diversity as the variance of community data: dissimilarity coefficients and partitioning. - Ecology Letters 16: 951–963.

Lerman, S. B., Larson, K. L., Narango, D. L., Goddard, M. A. and Marra, P. P. 2023. Humanity for Habitat: Residential Yards as an Opportunity for Biodiversity Conservation. - BioScience 73: 671–689.

Lin, B., Meyers, J. and Barnett, G. 2015. Understanding the potential loss and inequities of green space distribution with urban densification. - Urban Forestry & Urban Greening 14: 952–958.

Lososová, Z., Tichý, L., Divíšek, J., Čeplová, N., Danihelka, J., Dřevojan, P., Fajmon, K., Kalníková, V., Kalusová, V., Novák, P., Řehořek, V., Wirth, T. and Chytrý, M. 2018. Projecting potential future shifts in species composition of European urban plant communities. - Diversity and Distributions 24: 765–775.

Mack, R. N. and Lonsdale, W. M. 2001. Humans as Global Plant Dispersers: Getting More Than We Bargained For: Current introductions of species for aesthetic purposes present the largest single challenge for predicting which plant immigrants will become future pests. - BioScience 51: 95– 102.

Marchin, R. M., Esperon-Rodriguez, M., Tjoelker, M. G. and Ellsworth, D. S. 2022. Crown dieback and mortality of urban trees linked to heatwaves during extreme drought. - Science of The Total Environment 850: 157915.

Mata, L., Andersen, A. N., Morán-Ordóñez, A., Hahs, A. K., Backstrom, A., Ives, C. D., Bickel, D., Duncan, D., Palma, E., Thomas, F., Cranney, K., Walker, K., Shears, I., Semeraro, L., Malipatil, M., Moir, M. L., Plein, M., Porch, N., Vesk, P. A., Smith, T. R. and Lynch, Y. 2021. Indigenous plants promote insect biodiversity in urban greenspaces. - Ecological Applications 31: e02309.

Meineke, E. K. and Frank, S. D. 2018. Water availability drives urban tree growth responses to herbivory and warming. - Journal of Applied Ecology 55: 1701–1713.

Milano, M., Reynard, E., Köplin, N. and Weingartner, R. 2015. Climatic and anthropogenic changes in Western Switzerland: Impacts on water stress. - Science of The Total Environment 536: 12–24.

Misra, R. K., Patel, J. H. and Baxi, V. R. 2010. Reuse potential of laundry greywater for irrigation based on growth, water and nutrient use of tomato. - Journal of Hydrology 386: 95–102.

Ruile, S., Moretti, M., Ghazoul, J., Fournier, B., Grothe, V., Dädlow, L. and Abella, J. C. 2026. The human handprint in shaping plant diversity in urban environments. in press.

Russo, A., Esperon-Rodriguez, M., St-Denis, A. and Tjoelker, M. G. 2025. Native vs. Non-Native Plants: Public Preferences, Ecosystem Services, and Conservation Strategies for Climate-Resilient Urban Green Spaces. - Land in press.

Sikorski, P., Gawryszewska, B., Sikorska, D., Chormański, J., Schwerk, A., Jojczyk, A., Ciężkowski, W., Archiciński, P., Łepkowski, M., Dymitryszyn, I., Przybysz, A., Wińska-Krysiak, M., Zajdel, B., Matusiak, J. and Łaszkiewicz, E. 2021. The value of doing nothing – How informal green spaces can provide comparable ecosystem services to cultivated urban parks. - Ecosystem Services 50: 101339.

Simkin, R. D., Seto, K. C., McDonald, R. I. and Jetz, W. 2022. Biodiversity impacts and conservation implications of urban land expansion projected to 2050. - Proceedings of the National Academy of Sciences 119: e2117297119.

Speak, A. F., Mizgajski, A. and Borysiak, J. 2015. Allotment gardens and parks: Provision of ecosystem services with an emphasis on biodiversity. - Urban Forestry & Urban Greening 14: 772–781.

Spilková, J. and Vágner, J. 2016. The loss of land devoted to allotment gardening: The context of the contrasting pressures of urban planning, public and private interests in Prague, Czechia. - Land Use Policy 52: 232–239.

Swan, C. M., Pickett, S. T. A., Szlavecz, K., Warren, P. and Willey, K. T. 2011. Biodiversity and Community Composition in Urban Ecosystems: Coupled Human, Spatial, and Metacommunity Processes (JH Breuste, T Elmqvist, G Guntenspergen, P James, and NE McIntyre, Eds.). - Urban Ecology: 179–186.

Swiss Federal Statistical Office 2025. Household scenarios by canton and household size. - Federal Statistical Office (BFS).

Swiss Federal Statistical Office 2026. Swiss Land Use Statistics. - Federal Statistical Office (BFS).

Tartaglia, E. S. and Aronson, M. F. J. 2024. Plant native: comparing biodiversity benefits, ecosystem services provisioning, and plant performance of native and non-native plants in urban horticulture.- Urban Ecosyst 27: 2587–2611.

Thomas, C. D., Cameron, A., Green, R. E., Bakkenes, M., Beaumont, L. J., Collingham, Y. C., Erasmus, B. F. N., de Siqueira, M. F., Grainger, A., Hannah, L., Hughes, L., Huntley, B., van Jaarsveld, A. S., Midgley, G. F., Miles, L., Ortega-Huerta, M. A., Townsend Peterson, A., Phillips, O. L. and Williams, S. E. 2004. Extinction risk from climate change. - Nature 427: 145–148.

Threlfall, C. G., Ossola, A., Hahs, A. K., Williams, N. S. G., Wilson, L. and Livesley, S. J. 2016. Variation in Vegetation Structure and Composition across Urban Green Space Types. - Front. Ecol. Evol. in press.

Threlfall, C. G., Mata, L., Mackie, J. A., Hahs, A. K., Stork, N. E., Williams, N. S. G. and Livesley, S. J. 2017. Increasing biodiversity in urban green spaces through simple vegetation interventions. - Journal of Applied Ecology 54: 1874–1883.

Tomatis, F., Egerer, M., Correa-Guimaraes, A. and Navas-Gracia, L. M. 2023. Urban Gardening in a Changing Climate: A Review of Effects, Responses and Adaptation Capacities for Cities. - Agriculture in press.

Urban, M. C., Alberti, M., De Meester, L., Zhou, Y., Verrelli, B. C., Szulkin, M., Schmidt, C., Savage, A. M., Roberts, P., Rivkin, L. R., Palkovacs, E. P., Munshi-South, J., Malesis, A. N., Harris, N. C., Gotanda, K. M., Garroway, C. J., Diamond, S. E., Roches, S. D., Charmantier, A. and Brans, K. I. 2024. Interactions between climate change and urbanization will shape the future of biodiversity. - Nat. Clim. Chang.: 1–12.

Van der Veken, S., Hermy, M., Vellend, M., Knapen, A. and Verheyen, K. 2008. Garden plants get a head start on climate change. - Frontiers in Ecology and the Environment 6: 212–216.

Vanden Broeck, A., Cox, K., Melosik, I., Maes, B. and Smets, K. 2018. Genetic diversity loss and homogenization in urban trees: the case of Tilia × europaea in Belgium and the Netherlands. - Biodivers Conserv 27: 3777–3792.

Varquez, A. C. G. and Kanda, M. 2018. Global urban climatology: a meta-analysis of air temperature trends (1960–2009). - npj Clim Atmos Sci 1: 32.

Vega, K. A. and Küffer, C. 2021. Promoting wildflower biodiversity in dense and green cities: The important role of small vegetation patches. - Urban Forestry & Urban Greening 62: 127165.

Wagstaffe, B. and Dennis, M. 2025. Patch-level floral diversity and structural complexity drive differences in bumblebee richness and composition across different urban green space types. - Urban Ecosyst 28: 217.

WFO (2026): World Flora Online. Published on the Internet; http://www.worldfloraonline.org. Accessed on: 25 Sept 2025

Yin, Y., Liu, F., Manoli, G. and Cook, L. M. 2026. The cooling effect of urban green spaces in a future climate: projections for three Swiss cities. - Climate Services 43: 100700.

Young, C. and Bauer, N. 2022. Eviction from paradise: Lived experience, psycho-social and health effects of allotment garden loss. - Urban Forestry & Urban Greening 75: 127708.

Young, C., Hofmann, M., Frey, D., Moretti, M. and Bauer, N. 2020. Psychological restoration in urban gardens related to garden type, biodiversity and garden-related stress. - Landscape and Urban Planning 198: 103777.

Zhao, J., Ouyang, Z., Zheng, H., Zhou, W., Wang, X., Xu, W. and Ni, Y. 2010. Plant species composition in green spaces within the built-up areas of Beijing, China. - Plant Ecol 209: 189–204.

Zhao, L., Oleson, K., Bou-Zeid, E., Krayenhoff, E. S., Bray, A., Zhu, Q., Zheng, Z., Chen, C. and Oppenheimer, M. 2021. Global multi-model projections of local urban climates. - Nat. Clim. Chang. 11: 152–157.

