## Supplementary Material for "Joint effects of climate change and urbanization accelerate plant biodiversity loss across urban green spaces"

**Table of content**

Figure S1: Species accumulation curves for native and non-native plants across urban green space types and cities

Figure S2: Native and non-native plant species richness across urban green space types

Figure S3: Growth form composition of native and non-native plant communities across urban green space types

Figure S4: Local contributions to biodiversity (LCBD) of native and non-native plant communities across urban green space types

Figure S5: Differences in plant community composition across urban green space types

Figure S6: Projected vulnerability of native and non-native plant species to future climate change across urban green space types

Figure S7: Community-weighted thermal niche breadth of native and non-native plant species across urban green space types

Figure S8: Community-weighted thermal niche breadth across plant growth forms, species origins, and urban green space types

Figure S9: Differences in thermal niche characteristics and breadth between native and non-native plant species

Figure S10: Variation in thermal niche characteristics and breadth across plant growth forms and species origins

Figure S11: Historical trends and future projections of urban green space composition under alternative population growth scenarios

Figure S12: Projected gains and losses of native and non-native plant species under alternative urban growth scenarios

Figure S13: Projected changes in native and non-native plant species proportions under combined climate change and urban growth scenarios

Table S1: Pairwise comparisons of emission scenarios within urban green space types for native and non-native species

Table S2: Pairwise comparisons among urban green space types within emission scenarios for native and non-native species

Table S3: Projected changes in urban green space extent under alternative population growth scenarios, averaged across cities

Table S4: City-specific projections of urban green space extent under alternative population growth scenarios

Supplementary Material I: High-resolution urban microclimate modelling and projections under future climate scenarios

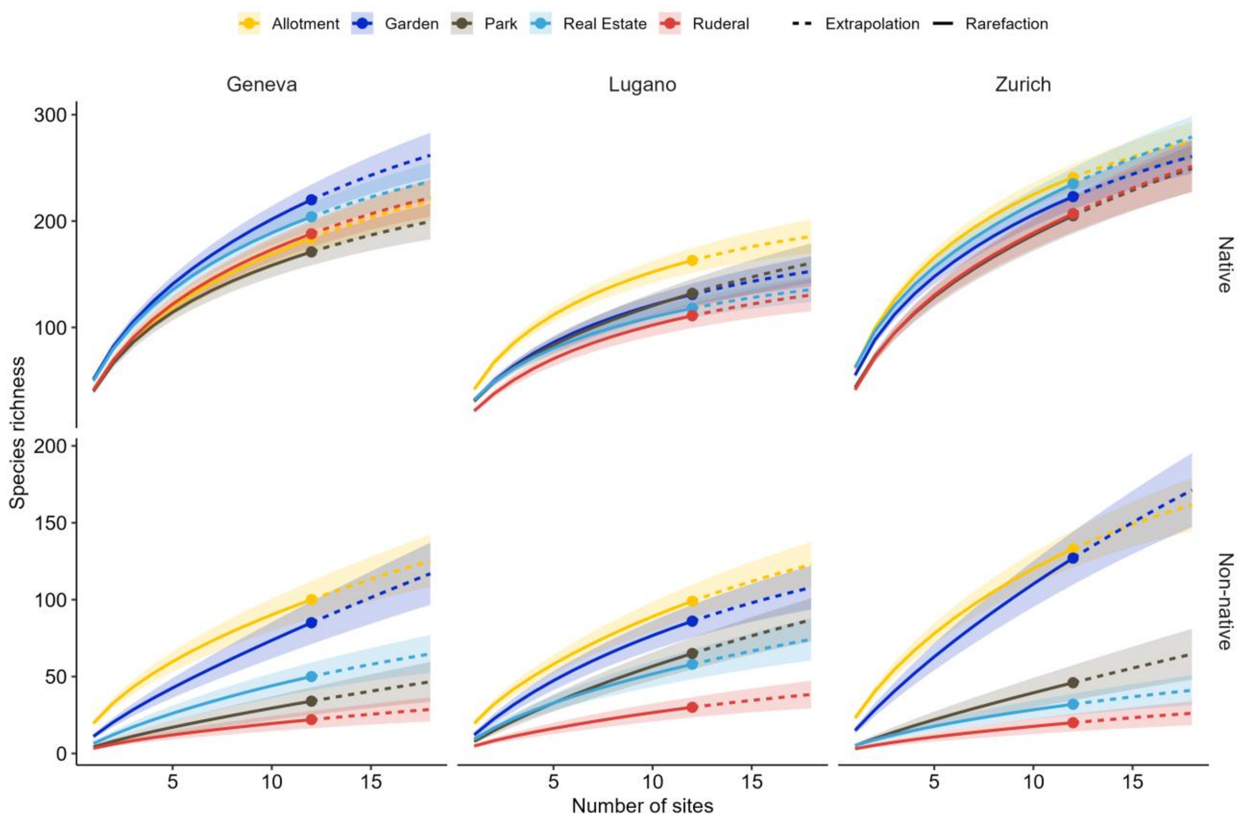


*Figure S1: Species accumulation curves for native (top) and non-native (bottom) plant species across UGS types in each study city. Solid lines represent interpolation (rarefaction) and dashed lines represent extrapolation of species richness with increasing numbers of sampled sites, with shaded ribbons showing 95% bootstrap confidence intervals. Points indicate the observed species richness.*

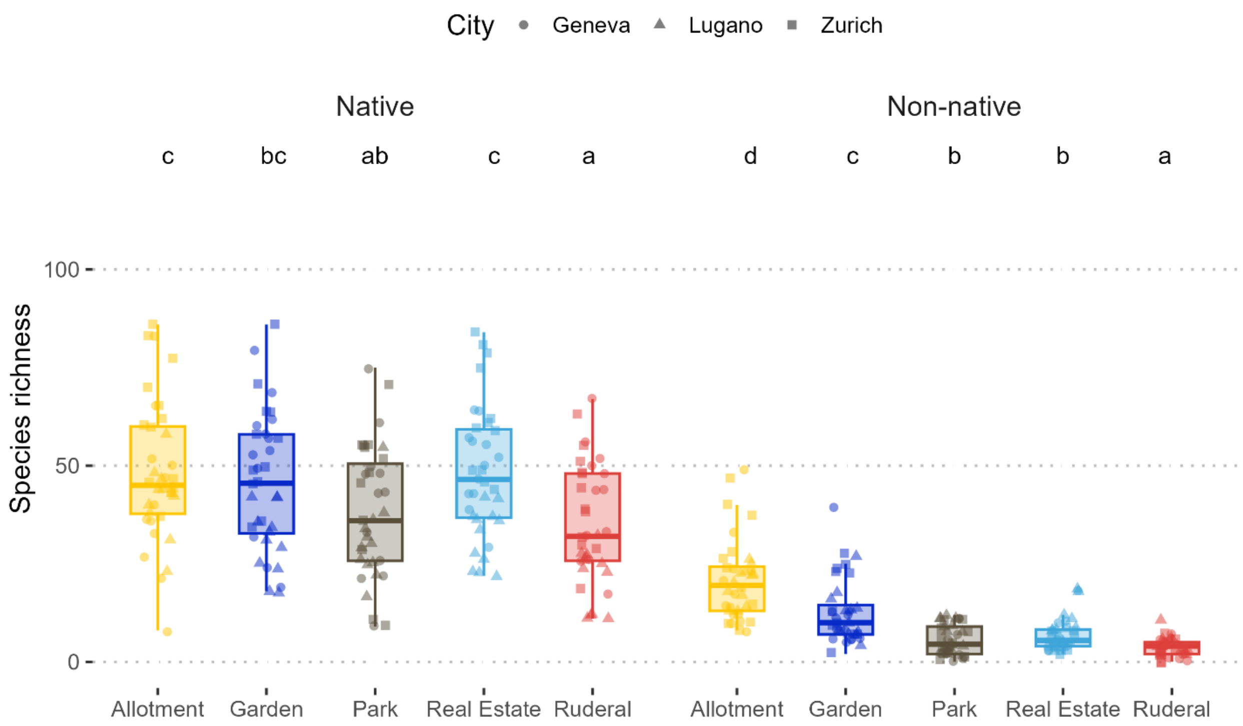


*Figure S2: Boxplot of native (left) and non-native (right) species richness per site across UGS type (color-coded). Shapes denote the different cities. Letters denote pairwise comparisons based on generalized linear mixed-effect models.*

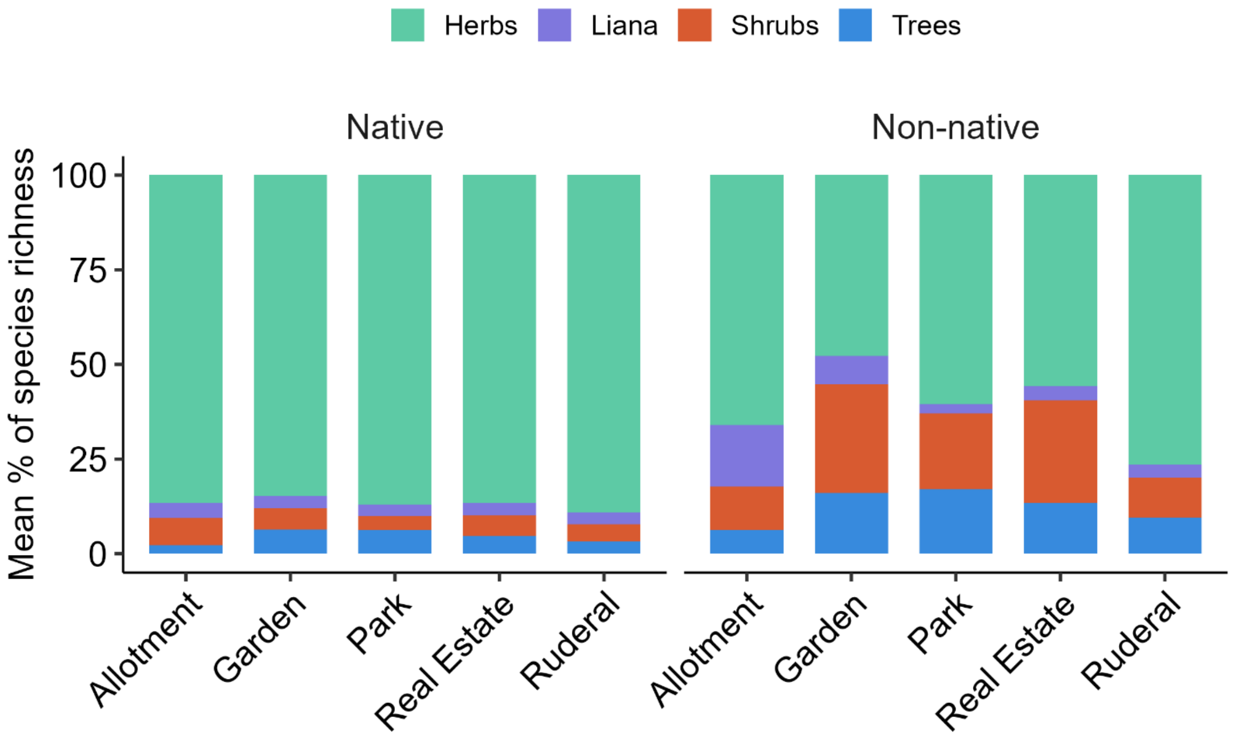


*Figure S3: Bar plots showing the average proportion of each growth form (herbs, lianas, shrubs, and trees; color-coded) per site across UGS types, for native (left) and non-native (right) species based on species' presence-absence.*

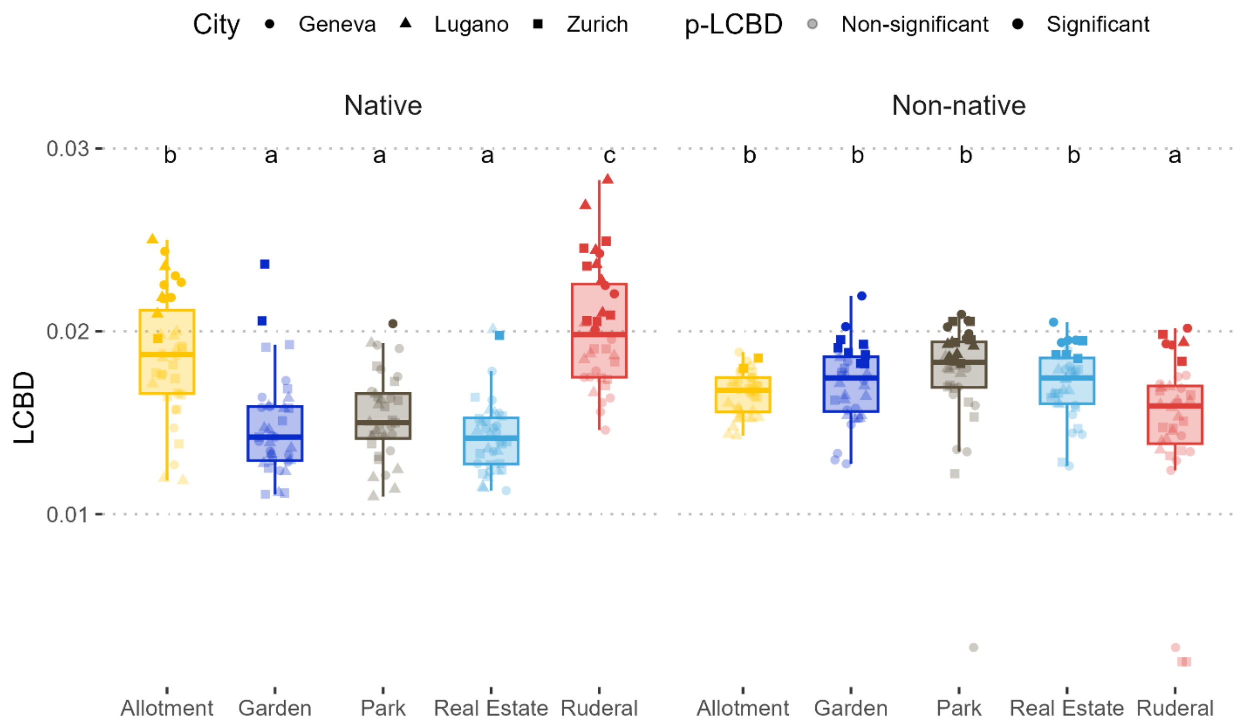


*Figure S4: Boxplot of native (left) and non-native (right) Local Contribution to Biodiversity (LCBD) per site across UGS type (color-coded). Shapes denote the different cities, and transparency denotes significant or non-significant contributions (based on 999 permutations). Letters denote pairwise comparisons based on linear mixed-effect models.*

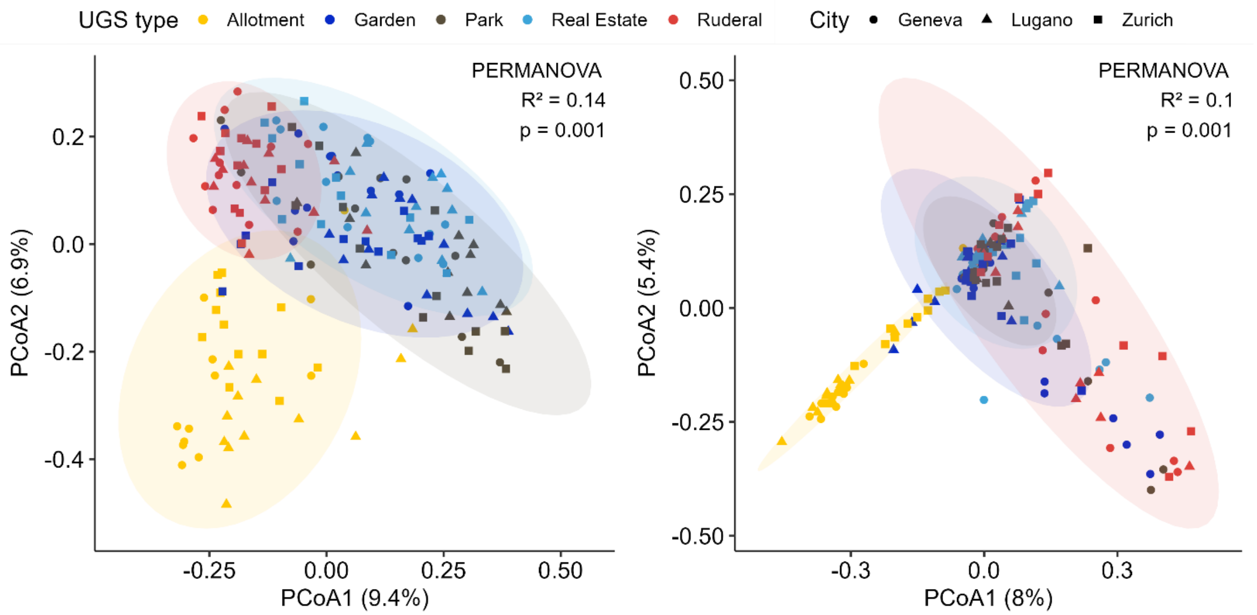


*Figure S5: Principal coordinate analysis (PCoA) showing compositional differences among sites across UGS types (color-coded), based on Bray–Curtis distances. Each point represents a site, with shape indicating city. Ellipses represent 95% confidence intervals. Results of a PERMANOVA testing compositional differences among UGS types are shown in the upper right corner.*

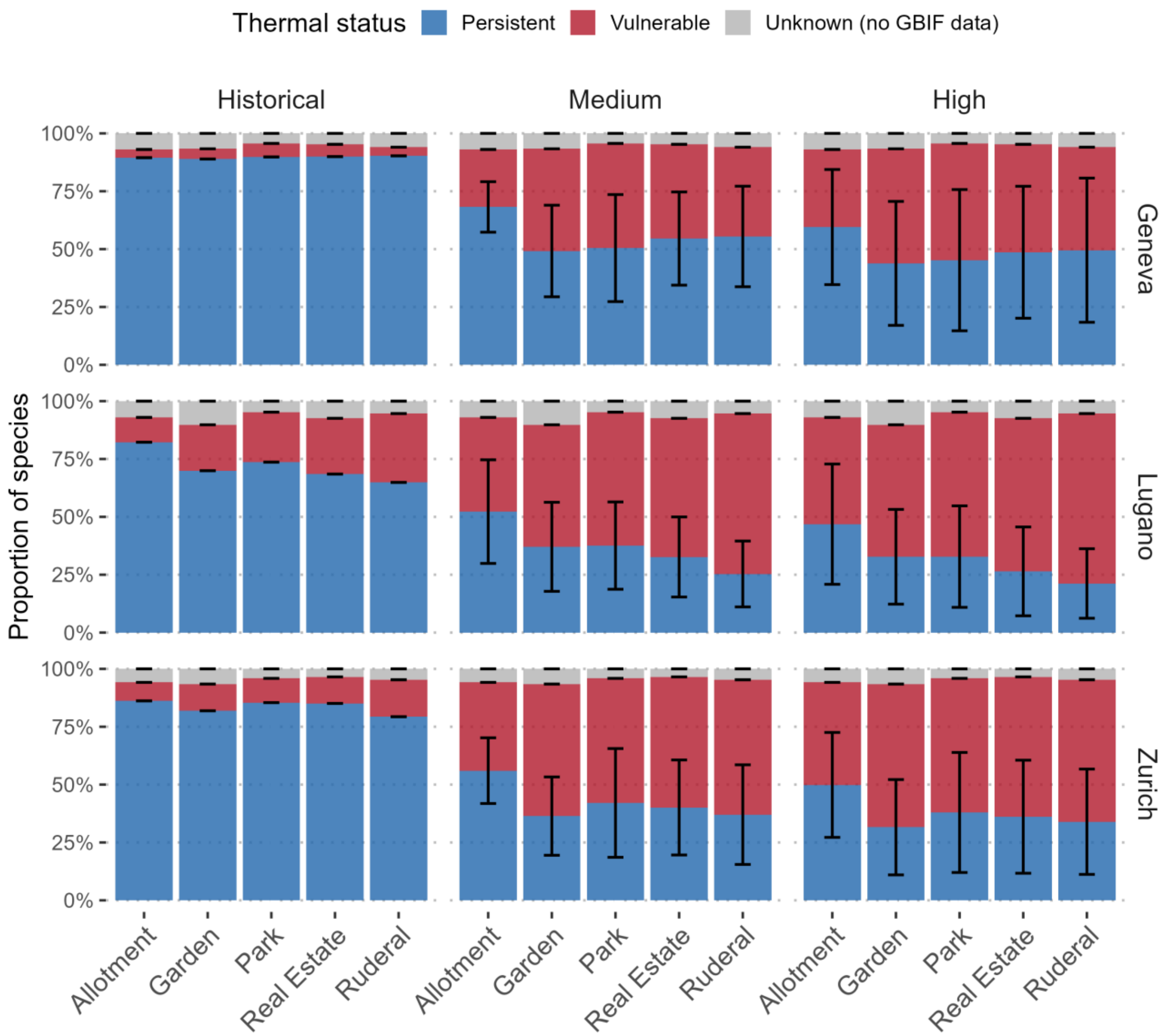


*Figure S6: Bar plots showing the proportion of species expected to remain within ("persistent"; blue) or exceed ("vulnerable"; red) their thermal limits, based on GBIF-derived climatic niches, under historical, medium-emission, and high-emission scenarios. Results are shown across the three cities (Geneva, Lugano, Zurich), climate models (GCMs MPI-ESM1-2-HR, UKESM1-0-LL, and GFDL-ESM4), and UGS types. Error bars indicate variation among climate model projections. Species with insufficient GBIF records to estimate climatic niches are classified as "unknown".*

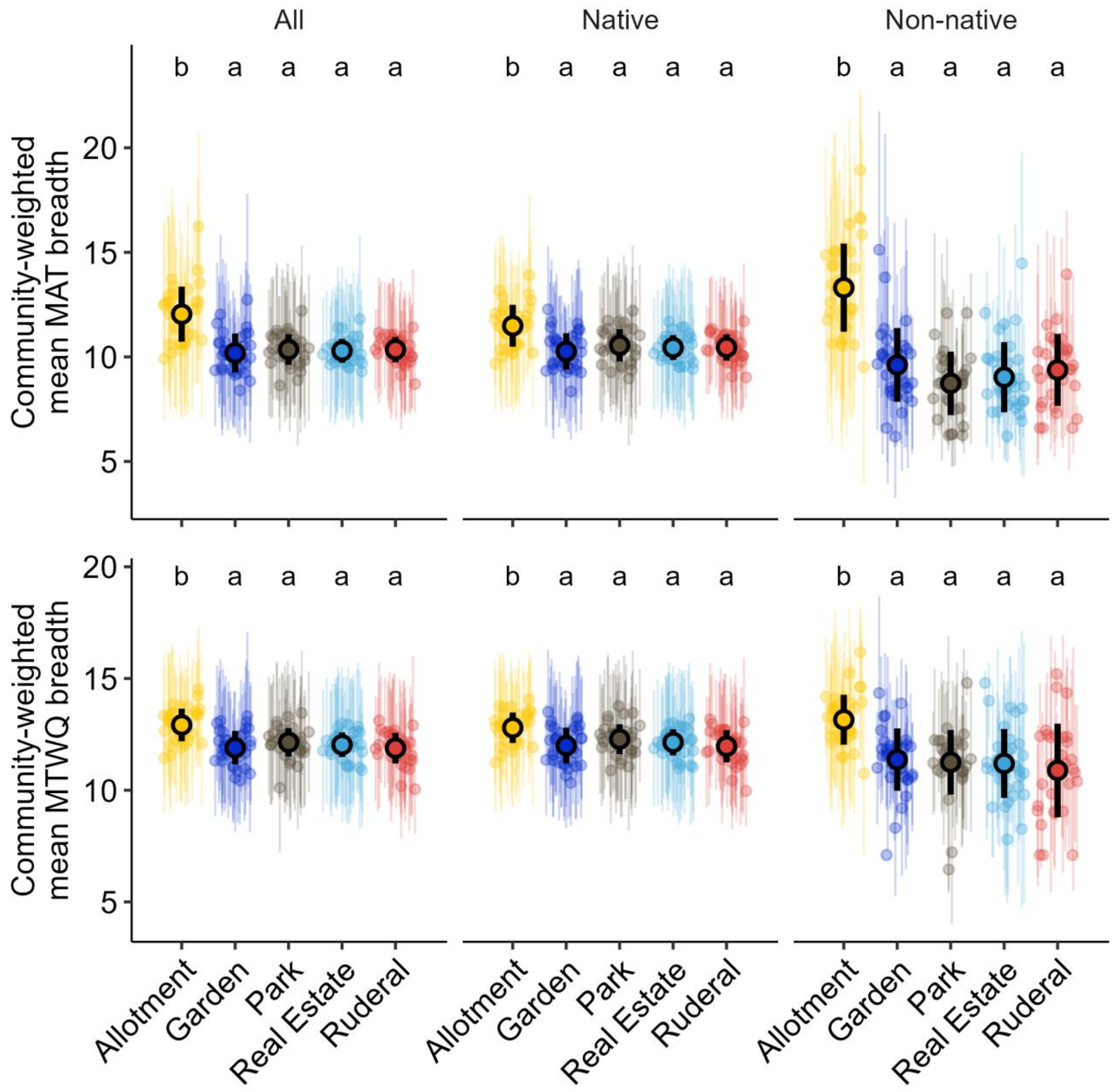


*Figure S7: Community-weighted thermal niche breadth across urban green space (UGS) types and species origins for mean annual temperature (MAT) breadth (top) and mean temperature of the warmest quarter (MTWQ) breadth (bottom). Points show site-level estimates, with error bars indicating standard deviation, Black points and ranges indicate means ± standard deviation for each UGS type. Letters indicate significant differences based on pairwise Wilcoxon tests with BH correction.*

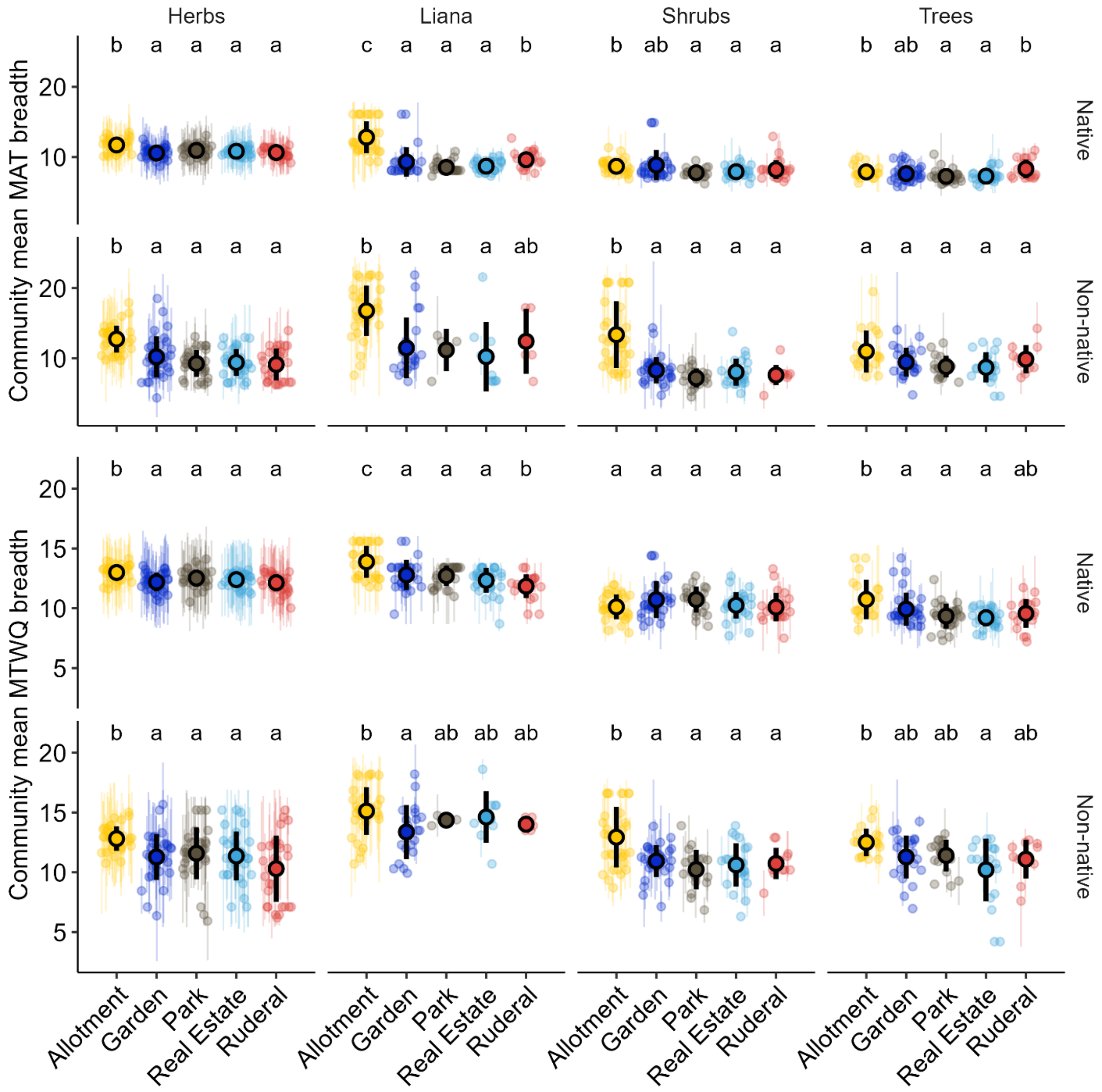


*Figure S8: Community-weighted thermal niche breadth across urban green space (UGS) types, growth form (herb, liana, shrubs, and trees), and origin (native, non-native) for mean annual temperature (MAT) breadth (top) and mean temperature of the warmest quarter (MTWQ) breadth (bottom). Points show site-level estimates, with error bars indicating standard deviation, Black points and ranges indicate means ± standard deviation for each UGS type. Letters indicate significant differences based on pairwise Wilcoxon tests with BH correction.*

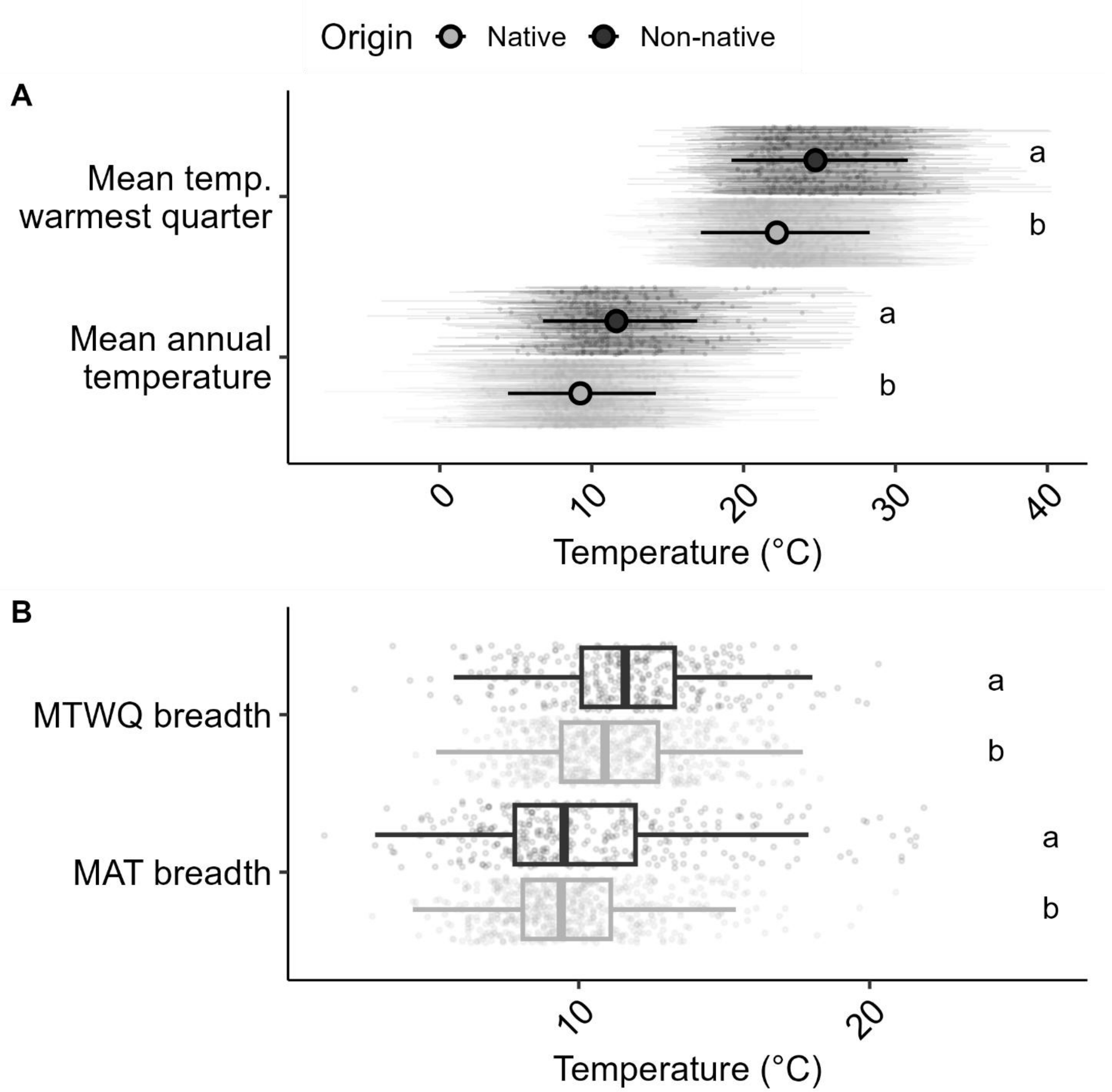


*Figure S9: Thermal niche characteristics of plant species across origin groups (native vs. non-native) for two climatic dimensions and their corresponding niche breadths. (A) Species-level thermal niches for mean annual temperature (MAT) and temperature of the warmest quarter (MTWQ), shown as points with thin error bars representing the 95% quantile range of observed occurrences. Black points and error bars indicate origin-level means with 95% confidence intervals. (B) Species-level thermal niche breadth, calculated as the difference between upper and lower bounds of each species' 95% occurrence-based thermal range, for MAT and MTWQ, displayed as boxplots with individual species values overlaid as points. Letters indicate significant differences between origins based on Tukey HSD post-hoc tests.*

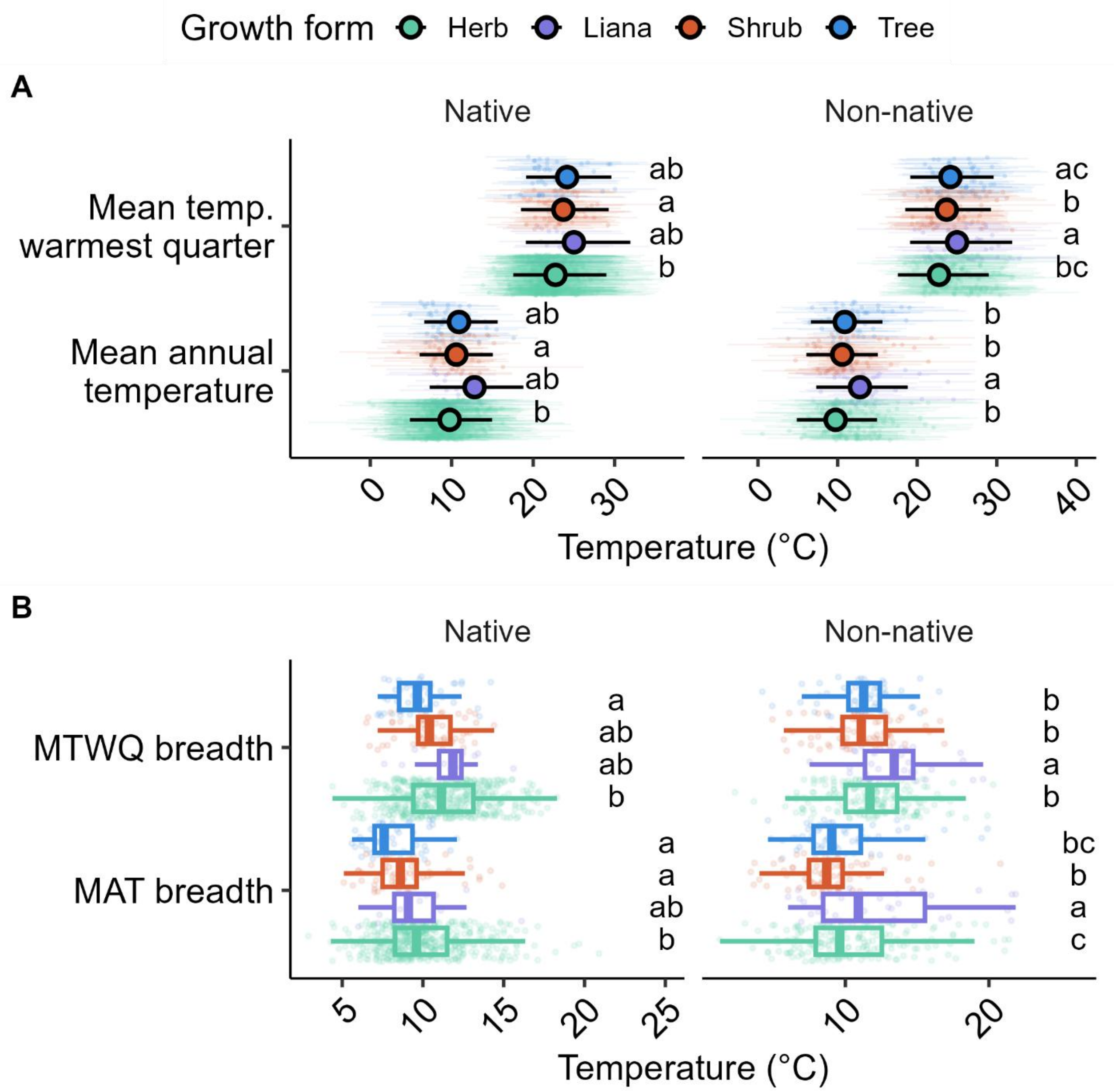


*Figure S10: Thermal niche characteristics of plant species across growth forms (herb, liana, shrubs, and trees) and origin groups (native vs. non-native) for two climatic dimensions and their corresponding niche breadths. (A) Species-level thermal niches for mean annual temperature (MAT) and temperature of the warmest quarter (MTWQ), shown as points with thin error bars representing the 95% quantile range of observed occurrences. Black points and error bars indicate origin-level means with 95% confidence intervals. (B) Species-level thermal niche breadth, calculated as the difference between upper and lower bounds of each species' 95% occurrence-based thermal range, for MAT and MTWQ, displayed as boxplots with individual species values*

*overlaid as points. Letters indicate significant differences between origins based on Tukey HSD post-hoc tests.*

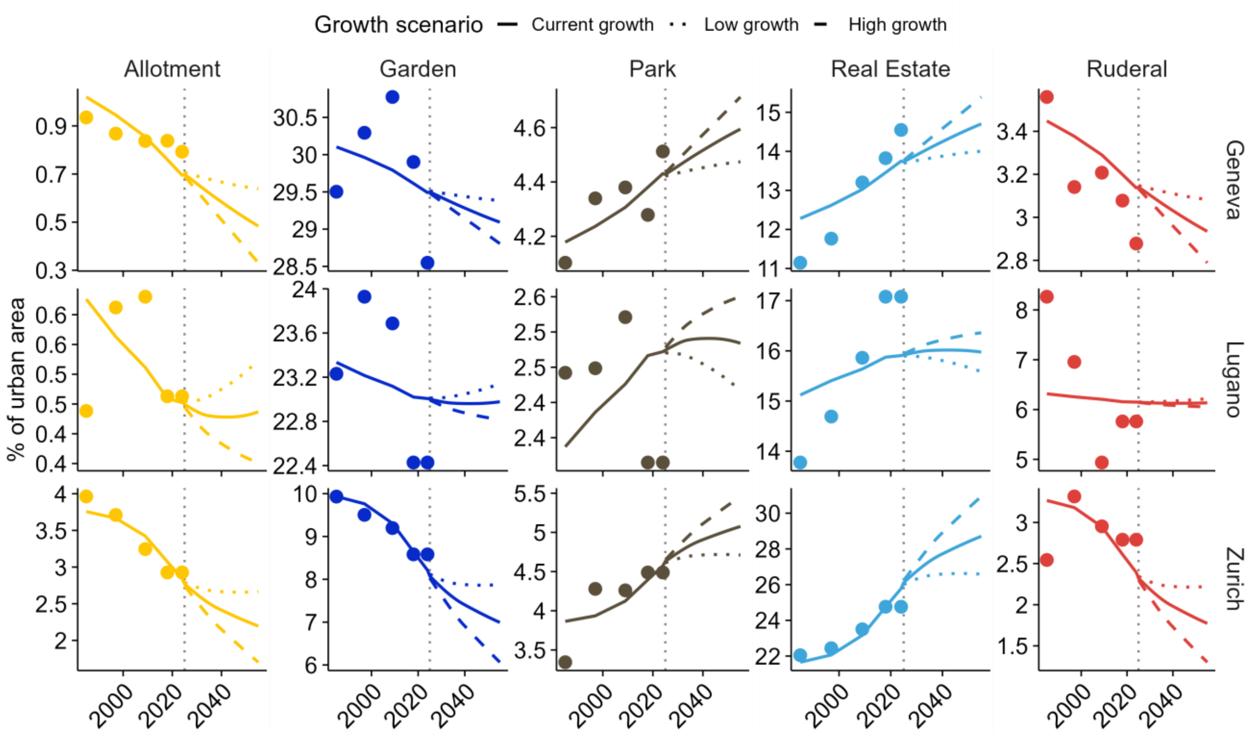


*Figure S11: Historical and projected trends in the proportion of UGS types (color-coded) as a function of human population growth for each city (Geneva, Lugano, Zurich). Points and dotted lines show observed historical values, while solid lines represent fitted model relationships. Projections beyond 2025 are shown under three population growth scenarios: current growth (solid), high growth (dashed), and low growth (dotted).*

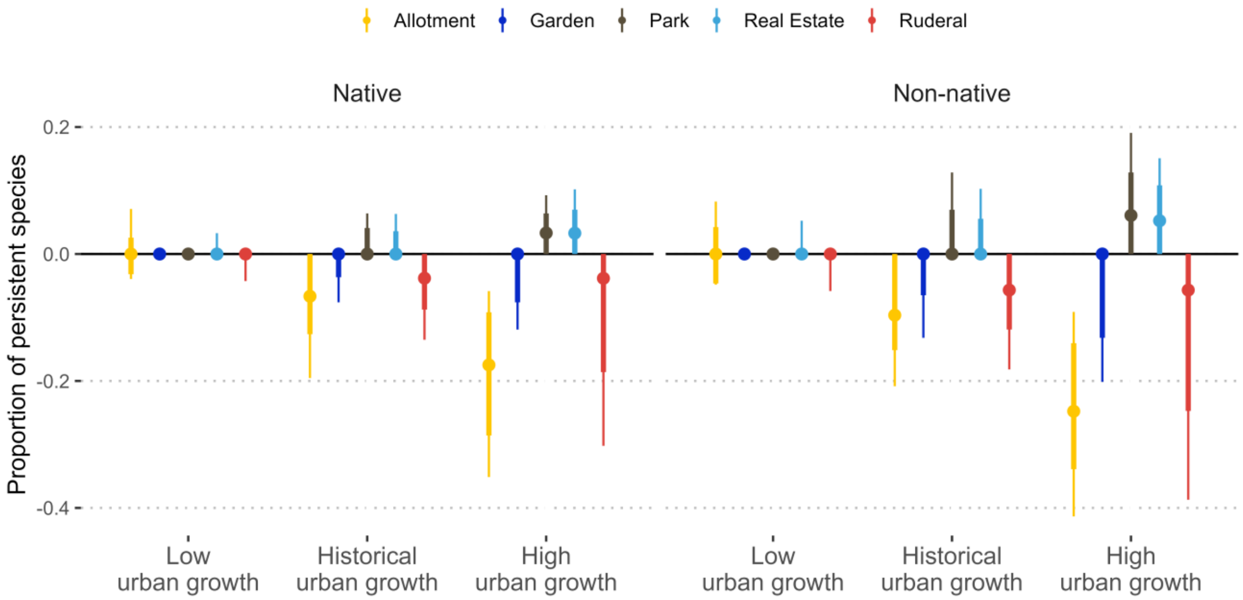


*Figure S12: Proportion of native (left) and non-native (right) species lost or gained through urbanization across urban green space (UGS) types (color-coded) under low, current, and high urban growth scenarios based on projected human population growth. Points indicate means, with thick and thin intervals representing the 50% and 75% quantile ranges, respectively. The horizontal black line indicates no net change; positive values represent species gains from site additions, whereas negative values represent species losses due to site removal. Results correspond to those shown in Figure 4 but are presented without climate scenarios for clarity.*

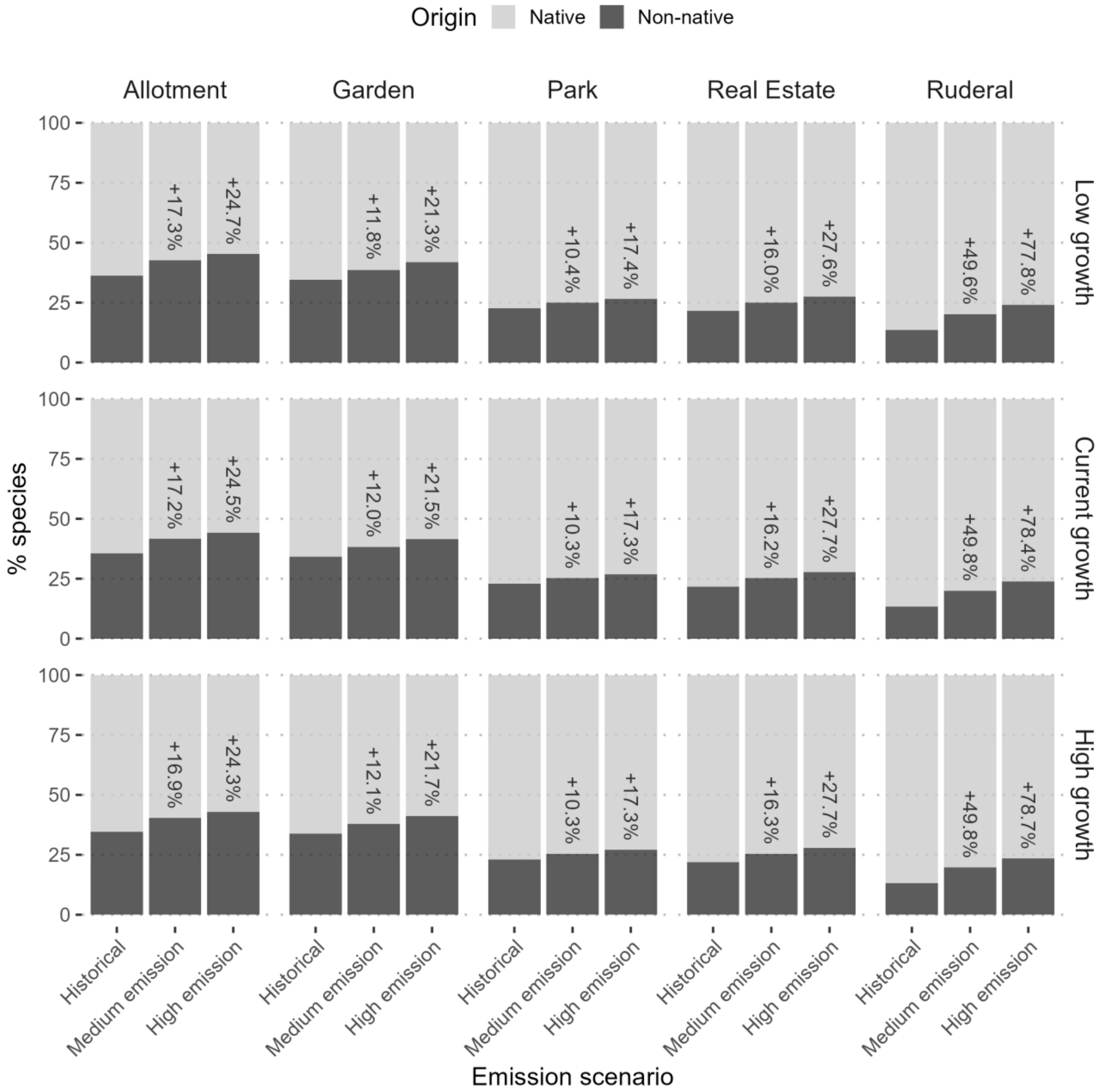


*Figure S13: Bar plots showing the proportion of native and non-native species under historical, medium-, and high-emission scenarios and urban growth scenarios (low, reference, high) across urban green space (UGS) types. Labels indicate the proportional increase in non-native species relative to current conditions.*

*Table S1: Pairwise comparison estimates, z-ratios, and p-values for contrasts among emission scenarios within each urban green space type, shown separately for native and non-native species.*

| *Origin* | *Urban green space type* | *Contrast* | *Estimate* | *z-ratio* | *p-value* |
|---|---|---|---|---|---|
| *Native* | *Allotment* | *Historical - Medium emission* | *-2.054* | *-2.051* | *0.100* |
| | | *Historical - High emission* | *-2.440* | *-2.436* | *0.039* |
| | | *Medium emission - High emission* | *-0.386* | *-5.959* | *<0.0001* |
| | *Garden* | *Historical - Medium emission* | *-2.429* | *-2.431* | *0.040* |
| | | *Historical - High emission* | *-2.692* | *-2.693* | *0.019* |
| | | *Medium emission - High emission* | *-0.262* | *-4.102* | *0.000* |
| | *Park* | *Historical - Medium emission* | *-2.250* | *-2.250* | *0.063* |
| | | *Historical - High emission* | *-2.504* | *-2.504* | *0.033* |
| | | *Medium emission - High emission* | *-0.254* | *-3.809* | *0.000* |
| | *Real Estate* | *Historical - Medium emission* | *-2.248* | *-2.250* | *0.063* |
| | | *Historical - High emission* | *-2.524* | *-2.526* | *0.031* |
| | | *Medium emission - High emission* | *-0.276* | *-4.453* | *<0.0001* |
| | *Ruderal* | *Historical - Medium emission* | *-2.170* | *-2.171* | *0.076* |
| | | *Historical - High emission* | *-2.408* | *-2.409* | *0.042* |
| | | *Medium emission - High emission* | *-0.238* | *-3.460* | *0.002* |
| *Non-native* | *Allotment* | *Historical - Medium emission* | *-1.281* | *-1.717* | *0.199* |
| | | *Historical - High emission* | *-1.630* | *-2.185* | *0.074* |
| | | *Medium emission - High emission* | *-0.349* | *-4.244* | *<0.0001* |
| | *Garden* | *Historical - Medium emission* | *-1.820* | *-2.416* | *0.042* |
| | | *Historical - High emission* | *-2.076* | *-2.756* | *0.016* |

| | | | | | |
|---|---|---|---|---|---|
| | | *Medium emission - High emission* | *-0.256* | *-2.556* | *0.029* |
| | *Park* | *Historical - Medium emission* | *-1.917* | *-2.484* | *0.035* |
| | | *Historical - High emission* | *-2.138* | *-2.772* | *0.015* |
| | | *Medium emission - High emission* | *-0.222* | *-1.594* | *0.248* |
| | *Real Estate* | *Historical - Medium emission* | *-1.678* | *-2.200* | *0.071* |
| | | *Historical - High emission* | *-1.973* | *-2.586* | *0.026* |
| | | *Medium emission - High emission* | *-0.295* | *-2.261* | *0.062* |
| | *Ruderal* | *Historical - Medium emission* | *-1.587* | *-2.027* | *0.106* |
| | | *Historical - High emission* | *-1.752* | *-2.238* | *0.065* |
| | | *Medium emission - High emission* | *-0.165* | *-0.972* | *0.595* |

131

*Table S2: Pairwise comparison estimates, z-ratios, and p-values for contrasts among urban green space type within each emission scenarios, shown separately for native and non-native species.*

| *Origin* | *Emission scenario* | *Contrast* | *Estimate* | *z-ratio* | *p-value* |
|---|---|---|---|---|---|
| *Native* | *Historical* | *Allotment - Garden* | *-0.55714* | *-3.496* | *0.0043* |
| | | *Allotment - Park* | *-0.57482* | *-3.538* | *0.0037* |
| | | *Allotment - Real Estate* | *-0.61518* | *-3.913* | *0.0009* |
| | | *Allotment - Ruderal* | *-0.84052* | *-5.269* | *<0.0001* |
| | | *Garden - Park* | *-0.01768* | *-0.119* | *1* |
| | | *Garden - Real Estate* | *-0.05804* | *-0.407* | *0.9942* |
| | | *Garden - Ruderal* | *-0.28338* | *-1.955* | *0.2883* |
| | | *Park - Real Estate* | *-0.04036* | *-0.277* | *0.9987* |
| | | *Park - Ruderal* | *-0.2657* | *-1.791* | *0.379* |
| | | *Real Estate - Ruderal* | *-0.22534* | *-1.581* | *0.5098* |
| | *Medium emission* | *Allotment - Garden* | *-0.93239* | *-14.148* | *<0.0001* |
| | | *Allotment - Park* | *-0.77058* | *-11.476* | *<0.0001* |
| | | *Allotment - Real Estate* | *-0.80903* | *-12.463* | *<0.0001* |
| | | *Allotment - Ruderal* | *-0.9566* | *-14.009* | *<0.0001* |
| | | *Garden - Park* | *0.161816* | *2.462* | *0.0994* |
| | | *Garden - Real Estate* | *0.123365* | *1.947* | *0.2924* |
| | | *Garden - Ruderal* | *-0.0242* | *-0.363* | *0.9963* |
| | | *Park - Real Estate* | *-0.03845* | *-0.594* | *0.9761* |
| | | *Park - Ruderal* | *-0.18602* | *-2.733* | *0.0493* |
| | | *Real Estate - Ruderal* | *-0.14757* | *-2.244* | *0.1638* |

| | | | | | |
|---|---|---|---|---|---|
| | *High emission* | *Allotment - Garden* | *-0.80863* | *-12.755* | *<0.0001* |
| | | *Allotment - Park* | *-0.63867* | *-9.871* | *<0.0001* |
| | | *Allotment - Real Estate* | *-0.69869* | *-11.226* | *<0.0001* |
| | | *Allotment - Ruderal* | *-0.80829* | *-12.248* | *<0.0001* |
| | | *Garden - Park* | *0.169957* | *2.61* | *0.0685* |
| | | *Garden - Real Estate* | *0.109941* | *1.756* | *0.3997* |
| | | *Garden - Ruderal* | *0.000336* | *0.005* | *1* |
| | | *Park - Real Estate* | *-0.06002* | *-0.937* | *0.8823* |
| | | *Park - Ruderal* | *-0.16962* | *-2.51* | *0.0884* |
| | | *Real Estate - Ruderal* | *-0.10961* | *-1.682* | *0.4453* |
| *Non-native* | *Historical* | *Allotment - Garden* | *-0.3176* | *-1.362* | *0.652* |
| | | *Allotment - Park* | *-0.404* | *-1.45* | *0.5954* |
| | | *Allotment - Real Estate* | *-0.5804* | *-2.266* | *0.1559* |
| | | *Allotment - Ruderal* | *-0.7099* | *-2.357* | *0.1273* |
| | | *Garden - Park* | *-0.0863* | *-0.294* | *0.9984* |
| | | *Garden - Real Estate* | *-0.2627* | *-0.963* | *0.8718* |
| | | *Garden - Ruderal* | *-0.3923* | *-1.244* | *0.7256* |
| | | *Park - Real Estate* | *-0.1764* | *-0.565* | *0.9801* |
| | | *Park - Ruderal* | *-0.306* | *-0.873* | *0.9067* |
| | | *Real Estate - Ruderal* | *-0.1295* | *-0.389* | *0.9951* |
| | *Medium emission* | *Allotment - Garden* | *-0.8561* | *-9.169* | *<0.0001* |
| | | *Allotment - Park* | *-1.0391* | *-8.995* | *<0.0001* |
| | | *Allotment - Real Estate* | *-0.9767* | *-8.834* | *<0.0001* |
| | | *Allotment - Ruderal* | *-1.0152* | *-7.531* | *<0.0001* |

| | | | | | |
|---|---|---|---|---|---|
| | | *Garden - Park* | *-0.1829* | *-1.504* | *0.5596* |
| | | *Garden - Real Estate* | *-0.1206* | *-1.033* | *0.8402* |
| | | *Garden - Ruderal* | *-0.1591* | *-1.136* | *0.7874* |
| | | *Park - Real Estate* | *0.0623* | *0.462* | *0.9907* |
| | | *Park - Ruderal* | *0.0239* | *0.154* | *0.9999* |
| | | *Real Estate - Ruderal* | *-0.0385* | *-0.254* | *0.9991* |
| | *High emission* | *Allotment - Garden* | *-0.763* | *-8.429* | *<0.0001* |
| | | *Allotment - Park* | *-0.912* | *-8.013* | *<0.0001* |
| | | *Allotment - Real Estate* | *-0.9229* | *-8.515* | *<0.0001* |
| | | *Allotment - Ruderal* | *-0.8315* | *-6.257* | *<0.0001* |
| | | *Garden - Park* | *-0.1491* | *-1.229* | *0.7345* |
| | | *Garden - Real Estate* | *-0.1599* | *-1.377* | *0.6427* |
| | | *Garden - Ruderal* | *-0.0686* | *-0.492* | *0.9881* |
| | | *Park - Real Estate* | *-0.0108* | *-0.08* | *1* |
| | | *Park - Ruderal* | *0.0805* | *0.518* | *0.9856* |
| | | *Real Estate - Ruderal* | *0.0913* | *0.603* | *0.9746* |

*Table S3: Projected land proportions of each UGS type across human population growth scenarios (current growth, high growth, and low growth), averaged across city.*

| UGS type | Urbanization scenario | % land (2050) | % land (2025) | % change |
|---|---|---|---|---|
| Allotment | Historical–medium emission | 1.039 | 1.306 | -20.471 |
| Allotment | High growth | 0.795 | 1.306 | -39.131 |
| Allotment | Low growth | 1.276 | 1.306 | -2.305 |
| Garden | Current growth | 19.688 | 20.192 | -2.494 |
| Garden | High growth | 19.229 | 20.192 | -4.768 |
| Garden | Low growth | 20.135 | 20.192 | -0.281 |
| Park | Current growth | 4.051 | 3.844 | 5.394 |
| Park | High growth | 4.240 | 3.844 | 10.310 |
| Park | Low growth | 3.867 | 3.844 | 0.607 |
| Real Estate | Current growth | 19.796 | 18.588 | 6.499 |
| Real Estate | High growth | 20.897 | 18.588 | 12.423 |
| Real Estate | Low growth | 18.724 | 18.588 | 0.732 |
| Ruderal | Current growth | 3.614 | 3.870 | -6.612 |
| Ruderal | High growth | 3.381 | 3.870 | -12.639 |
| Ruderal | Low growth | 3.841 | 3.870 | -0.744 |

*Table S4: Projected land proportions of each UGS type across human population growth scenarios (current growth, high growth, and low growth), averaged across city.*

| City | UGS type | Urbanization scenario | % land (2050) | % land (2025) | % change |
|---|---|---|---|---|---|
| Geneva | Allotment | Current growth | 0.484 | 0.699 | -30.771 |
| Zurich | Allotment | Current growth | 2.195 | 2.770 | -20.739 |
| Lugano | Allotment | Current growth | 0.437 | 0.450 | -2.813 |
| Lugano | Allotment | High growth | 0.352 | 0.450 | -21.764 |
| Zurich | Allotment | High growth | 1.702 | 2.770 | -38.553 |
| Geneva | Allotment | High growth | 0.331 | 0.699 | -52.600 |
| Geneva | Allotment | Low growth | 0.639 | 0.699 | -8.557 |
| Lugano | Allotment | Low growth | 0.523 | 0.450 | 16.406 |
| Zurich | Allotment | Low growth | 2.666 | 2.770 | -3.765 |
| Geneva | Garden | Current growth | 29.093 | 29.498 | -1.373 |
| Zurich | Garden | Current growth | 6.993 | 8.075 | -13.401 |
| Lugano | Garden | Current growth | 22.978 | 23.002 | -0.104 |
| Lugano | Garden | High growth | 22.817 | 23.002 | -0.801 |
| Zurich | Garden | High growth | 6.064 | 8.075 | -24.912 |
| Geneva | Garden | High growth | 28.806 | 29.498 | -2.347 |
| Geneva | Garden | Low growth | 29.386 | 29.498 | -0.382 |
| Lugano | Garden | Low growth | 23.141 | 23.002 | 0.604 |
| Zurich | Garden | Low growth | 7.879 | 8.075 | -2.433 |
| Geneva | Park | Current growth | 4.594 | 4.428 | 3.766 |
| Zurich | Park | Current growth | 5.076 | 4.630 | 9.621 |
| Lugano | Park | Current growth | 2.484 | 2.474 | 0.396 |

| | | | | | |
|---|---|---|---|---|---|
| Lugano | Park | High growth | 2.550 | 2.474 | 3.067 |
| Zurich | Park | High growth | 5.458 | 4.630 | 17.885 |
| Geneva | Park | High growth | 4.713 | 4.428 | 6.437 |
| Geneva | Park | Low growth | 4.474 | 4.428 | 1.047 |
| Lugano | Park | Low growth | 2.417 | 2.474 | -2.312 |
| Zurich | Park | Low growth | 4.711 | 4.630 | 1.747 |
| Geneva | Real Estate | Current growth | 14.704 | 13.732 | 7.074 |
| Zurich | Real Estate | Current growth | 28.708 | 26.113 | 9.939 |
| Lugano | Real Estate | Current growth | 15.975 | 15.918 | 0.359 |
| Lugano | Real Estate | High growth | 16.360 | 15.918 | 2.777 |
| Zurich | Real Estate | High growth | 30.937 | 26.113 | 18.476 |
| Geneva | Real Estate | High growth | 15.393 | 13.732 | 12.092 |
| Geneva | Real Estate | Low growth | 14.003 | 13.732 | 1.967 |
| Lugano | Real Estate | Low growth | 15.585 | 15.918 | -2.094 |
| Zurich | Real Estate | Low growth | 26.584 | 26.113 | 1.804 |
| Geneva | Ruderal | Current growth | 2.935 | 3.141 | -6.552 |
| Zurich | Ruderal | Current growth | 1.773 | 2.323 | -23.670 |
| Lugano | Ruderal | Current growth | 6.135 | 6.147 | -0.197 |
| Lugano | Ruderal | High growth | 6.054 | 6.147 | -1.524 |
| Zurich | Ruderal | High growth | 1.301 | 2.323 | -44.000 |
| Geneva | Ruderal | High growth | 2.789 | 3.141 | -11.201 |
| Geneva | Ruderal | Low growth | 3.083 | 3.141 | -1.822 |
| Lugano | Ruderal | Low growth | 6.218 | 6.147 | 1.148 |
| Zurich | Ruderal | Low growth | 2.223 | 2.323 | -4.297 |

### *Supplementary Material I*

Following the framework developed by Dietzel et al. (2025), we compiled high-resolution air temperature observations from 91 urban weather stations distributed across Zurich, capturing a broad range of climatic and environmental conditions. Local temperature was recorded at 15-minute intervals at 2 m above ground over four years (January 2020–December 2023) and subjected to extensive quality control, including removal of outliers and flatliners and correction for sensor heat accumulation. From these observations, temperature-based bioclimatic variables (bio1–bio11) were derived. In parallel, we obtained CHELSA V2 bioclimatic projections (30 arc seconds), covering historical (1981–2010) and mid-century (2041–2070) periods, two shared socio-economic pathways (SSP3-7.0 and SSP5-8.5), and five global circulation models (GFDL-ESM4, IPSL-CM6A-LR, MPI-ESM1-2-HR, MRI-ESM2-0, UKESM1-0-LL), capturing a broad range of climatic and socio-economic uncertainty. Although both SSP scenarios represent relatively high emissions pathways, we refer to them as “medium” and “high” for clarity.

The original framework of Dietzel et al. (2025) accounted for urban overwarming by estimating empirical deviations ($\Delta k$) in bioclimatic conditions for each station relative to rural reference conditions, thereby quantifying how local environmental conditions modify regional climate signals without replacing the CHELSA baseline. However, because these deviations are defined relative to a single reference station, the original formulation is not directly transferable across cities. To obtain a transferable approach and enable projections for Geneva and Lugano, we therefore adapted the framework by calculating deviations for each bioclimatic variable k at station i relative to the mean across all stations ($\Delta k = ki - \bar{k}$). This formulation provides a consistent definition of urban–rural contrasts that is independent of any single location and allows spatial offsets to be estimated across heterogeneous urban environments. To obtain city-wide high-resolution projections of microclimatic conditions for each city, bioclimatic variable and climate scenario, we first calculated the mean city-wide offset by adding the predicted Δk at the rural reference weather station to its projected (CHELSA) bioclimatic condition, and then added this mean city-wide offset to the predicted Δk in each grid cell.

Covariates used to model Δk were selected based on their availability across all three cities and their ability to capture local and landscape-scale variation in urban structure, including topography, land use, and vegetation. Predictor sets were refined to better represent urban green structure, with particular emphasis on NDVI and canopy height as proxies for vegetation cover and land-use intensity. Additional predictors included elevation, slope, aspect, hillshade, aggregated NDVI at local and 100 m resolution (Külling et al. 2024), and grey surface fractions at multiple spatial scales (local, 100 m, 200 m, and 500 m) (Price et al. 2021). Model configuration, including parameter settings, otherwise followed Dietzel et al. (2025) and remained identical to the original specification; an advanced predictor set was not considered in our study.

Price, B., Kolecka, N., Huber, N., Rüetschi, M., Nussbaumer, A., & Ginzler, C. (2024). The Habitat Map of Switzerland v1_1 2024. EnviDat. https://www.envidat.ch/dataset/9b4526f9-ef08-4f11-860f-788287fceaec

Dietzel, A., Moretti, M., Perrelet, K., Cook, L.M., 2025. Urban heat exacerbates climatic risks to urban biodiversity. Npj Urban Sustain. 6, 4. https://doi.org/10.1038/s42949-025-00309-6

Külling, N., Adde, A., Fopp, F., Schweiger, A.K., Broennimann, O., Rey, P.-L., Giuliani, G., Goicolea, T., Petitpierre, B., Zimmermann, N.E., Pellissier, L., Altermatt, F., Lehmann, A., Guisan, A., 2024. SWECO25: a cross-thematic raster database for ecological research in Switzerland. Sci. Data 11, 21. https://doi.org/10.1038/s41597-023-02899-1
